# Monoclonal antibodies targeting surface exposed epitopes of *Candida albicans* cell wall proteins confer *in vivo* protection in an infection model

**DOI:** 10.1101/2021.09.29.462385

**Authors:** Soumya Palliyil, Mark Mawer, Sami A Alawfi, Lily Fogg, Tyng H Tan, Giuseppe Buda De Cesare, Louise A Walker, Donna M MacCallum, Andrew J Porter, Carol A Munro

## Abstract

MAb based immunotherapies targeting systemic and deep-seated fungal infections are still in their early stages of development with currently no licensed antifungal mAbs available for patients at risk. The cell wall glycoproteins of *Candida albicans* are of particular interest as potential targets for therapeutic antibody generation due to their extracellular location and key involvement in fungal pathogenesis. Here we describe the generation of recombinant human antibodies specifically targeting two key cell wall proteins (CWPs) in *C. albicans* - Utr2 and Pga31. These antibodies were isolated from a phage display antibody library using peptide antigens representing the surface exposed regions of CWPs expressed at elevated levels during *in vivo* infection. Reformatted human-mouse chimeric mAbs preferentially recognised *C. albicans* hyphal forms compared to yeast cells and an increased binding was observed when the cells were grown in the presence of the antifungal agent caspofungin. In J774.1 macrophage interaction assays, mAb pre-treatment resulted in a faster engulfment of *C. albicans* cells suggesting a role of the CWP antibodies as opsonising agents during phagocyte recruitment. Finally, in a series of clinically predictive, mouse models of systemic candidiasis, our lead mAb achieved an improved survival (83%) and several log reduction of fungal burden in the kidneys, similar to levels achieved for the fungicidal drug caspofungin, and superior to any anti-*Candida* mAb therapeutic efficacy reported to date.

## INTRODUCTION

Invasive fungal infections (IFIs) are serious, life-threatening conditions typically affecting individuals with a compromised immune system including patients with haematological malignancies, those undergoing cytotoxic chemotherapy and organ transplantation (1). Ironically, due to advancements in immunomodulatory drugs (including antibodies), patients suffering from cancer and other complex health conditions are often left with a temporary but weakened immune system due to their medication. This drug-induced immune suppression has caused a significant rise in life-threatening systemic and organ specific fungal infections (1). Clinicians suffer the frustration of successfully controlling cancers only to see their patients succumb to these often hard to treat infections. Opportunistic pathogenic fungi, including members of *Aspergillus*, *Candida* and *Cryptococcus* species, are responsible for invasive fungal infections and the death of at least 1.7 million people each year globally (2). Population-based surveillance studies show that the yearly incidence of invasive *Candida albicans* infections, and related species including *Candida parapsilosis, Candida glabrata* and *Candida tropicalis,* can be as high as 21 per 100,000 in some geographies (1). Furthermore, the ongoing global COVID-19 pandemic has fuelled an increase in secondary fungal superinfections, such as SARS-CoV-2 associated pulmonary aspergillosis (CAPA), with mortality rates greater than 40% in almost all study cohorts (3).

Currently, IFIs are treated with antifungal agents belonging to four main drug classes, and include amphotericin B, fluconazole, voriconazole, caspofungin and 5-flucytosine. The recalcitrance of some infections has encouraged longer term treatment regimens and even their prophylactic use for some surgeries and organ transplantation. This unmanaged use of a limited drug armory has inevitably led to the emergence of antifungal drug resistance in many fungal genera (4) (5). Certain *Candida* species, such as *Candida auris,* are of particular concern in several countries with some isolates showing reduced susceptibility to fluconazole, amphotericin B and echinocandins, with clinicians labelling *C. auris* the MRSA of the fungal world (6) (7). The increasingly well-documented shortcomings of our existing antifungals (toxicity, complex drug-drug interactions, emergence of multidrug resistance strains) and the intrinsic ability of certain fungal species to evade drug therapies has accelerated the need to develop novel “first-in-class” alternatives to tackle these life-threatening conditions.

In all pathogenic fungi of consequence, the cell wall is a dynamic structure continuously changing in response to body/culture conditions and environmental stimuli. The cell wall of *C. albicans* is covered in an outer layer of glycoproteins that play important roles in pathogenesis and mediating interactions between the host and the fungus (8). These proteins “immune-mask” fungal β-glucans from recognition by the mammalian β-glucan receptor dectin-1 (9). Some of these cell surface glycoproteins, including adhesins, invasins and superoxide dismutases, are also important virulence factors in their own right. Often shed by the invading fungus, these proteins promote adherence of *C. albicans* to host cells, mediate tissue invasion and combat oxidative burst defences (10). Many cell surface glycoproteins are post-translationally modified by the addition of Glycosylphosphatidylinositol (GPI)-anchor and can be either fungal plasma membrane localised or translocated into the cell wall, where they are covalently attached to the β-(1, 6)-glucan polymer (11). More than 100 putative GPI-anchored proteins have been annotated in *C. albicans* using *in silico* analysis (12) (13), with some having enzymatic functions associated with cell wall biosynthesis and cell wall remodeling. Several studies have reported alterations in cell wall protein (CWP) composition and expression in response to changes in growth conditions including carbon source, iron limitation, hypoxia or antifungal drug challenge, indicating the possibility of up/down regulation of these proteins during *in vivo* infection (14). Whilst CWPs define the success of many fungal pathogens, some may also provide an “Achilles’ heel” which can be exploited in therapy. Neutralising antibodies against CWPs have been detected in patients’ sera and therefore represent an important source of antigens/epitopes for vaccine generation and therapeutic antibody development (15). GPI-anchored CWPs, such as Pga31 and Utr2, play important roles in cell wall integrity and assembly (16). Utr2 carries a glycoside hydrolase family 16 domain, predictive of transglycosidase activity that catalyses cross-links between β-(1,3)-glucan and chitin, as shown for the *S. cerevisiae* orthologue Crh1, and is involved in cell wall remodelling and maintenance (17). In *C. albicans, UTR2, CRH11,* and *CRH12* belong to the *CRH* gene family and are strongly regulated by calcineurin, a serine/threonine protein phosphatase involved in cell wall morphogenesis and virulence (18) (19). Mutants lacking *UTR2* exhibit defective cell wall organisation (inducing the cell integrity MAP kinase signalling pathway), reduction in adherence to mammalian cells and reduced virulence, resulting in the prolonged survival of animals in *in vivo* models of systemic infection (18) (20). Immunofluorescence staining locates Utr2 predominantly to the budding site of mother yeast cells, eventually forming a ring at the base of the neck, whereas during hyphal elongation, the protein is detected at the tip of the germ tube and as a ring at the septum (18). Utr2 co-localises to chitin-rich regions in yeast, pseudohyphal and hyphal forms (18). Another GPI-anchored glycoprotein Pga31, which has unknown function, is upregulated in the opaque form of *C. albicans* and after exposure to caspofungin (12) (21). A *pga31* null mutant exhibits decreased chitin content compared to the wildtype strain and increased sensitivity to caspofungin and cell wall perturbing agents such as Calcofluor white (CFW) and SDS (12). The low chitin phenotype of the *pga31* null mutant points to a role for Pga31 linked to chitin assembly during cell wall biogenesis and the maintenance of wall integrity under cellular stress. Given their established roles in cell wall remodelling, upregulation after caspofungin treatment (22) and enhanced expression in *in vivo* models of systemic candidiasis (23), we investigated the potential of Utr2 and Pga31 as therapeutic targets for the development of monoclonal antibodies (mAbs) to treat life-threatening fungal infections. MAb based therapies have seen unprecedented levels of success in cancer and autoimmune disorders, producing several blockbuster drugs including Humira®. This molecule class has also expanded into novel therapeutic modalities such as bispecific antibodies and antibody-drug conjugates (ADCs) (24) (25).

Cancer treatments were once dominated by toxic small molecule therapies requiring a balancing act between killing the cancer cells and damaging healthy cells. However, cancer therapy was revolutionised by targeted mAb therapies which have increased efficacy and reduced side-effects. The balancing act is also practiced by infectious disease clinicians treating life-threatening systemic fungal infections where toxic molecules are used to kill the fungus without “killing” the patient. Unfortunately, mAb technology has failed to generate significant impact in the infectious diseases field to date, but the increasing emergence of drug resistant fungal strains is accelerating the need for new therapeutic modalities. Currently, only a handful of antifungal mAbs are reported to show modest efficacy against *in vivo* infection and none of these have so far progressed into the clinic (26) (27) (28)

In this study, the *C. albicans* cell wall proteome was interrogated using trypsin digestion followed by LC-MS/MS analysis to identify several covalently linked CWPs, including Utr2, Pga31, and their surface exposed epitopes. These epitopes were used to generate monoclonal antibodies from a naïve human phage display antibody library. The ability of these recombinant mAbs to bind several pathogenic fungi was investigated with some showing fungal-specific, and others fungal species-specific binding. However, most importantly, their protective efficacy has been demonstrated in a murine model of systemic candidiasis, with a potency approaching that of more traditional antifungal drug classes.

## RESULTS

### Antigen design for recombinant antibody generation

Guided by the cell wall proteome analysis of various caspofungin susceptible and resistant strains of *C. albicans*, peptide sequences accessible to trypsin digestion were identified and matched with their respective fungal cell wall proteins (Tables 1A and 1B) (22). Based on the predicted β turn structures (algorithm NetTurnP 1.0) and hydropathy of these surface exposed regions, peptide sequences were selected as antigens representing the CWPs Utr2 and Pga31. β turn regions are often solvent exposed secondary structures and tend to have a relatively higher propensity for antibody binding (29). A small panel of trypsin-susceptible peptides from both proteins were custom synthesised and C terminally biotinylated via an additionally introduced lysine residue and these conjugates used as antigens for biopanning experiments.

**Table 1A.**
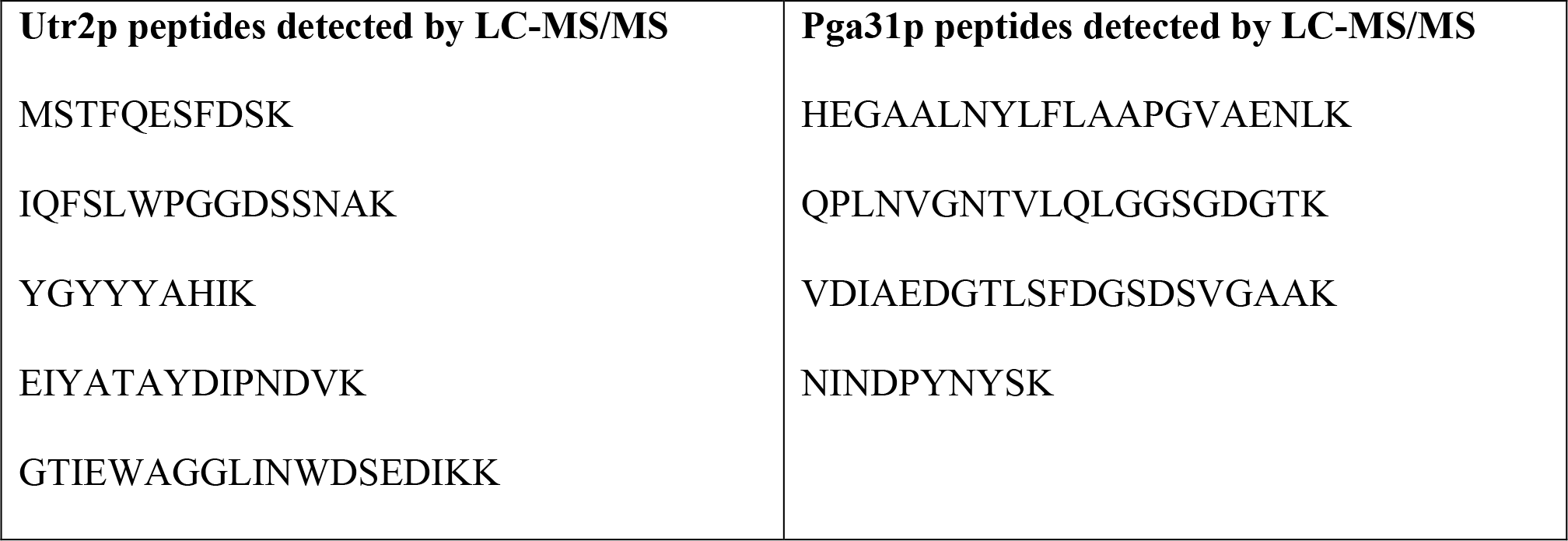
Amino acid sequences of the tryptic digested peptides identified in the cell wall proteome analysis of C. albicans SC5314 using LC-MS/MS method.

**Table 1B.**
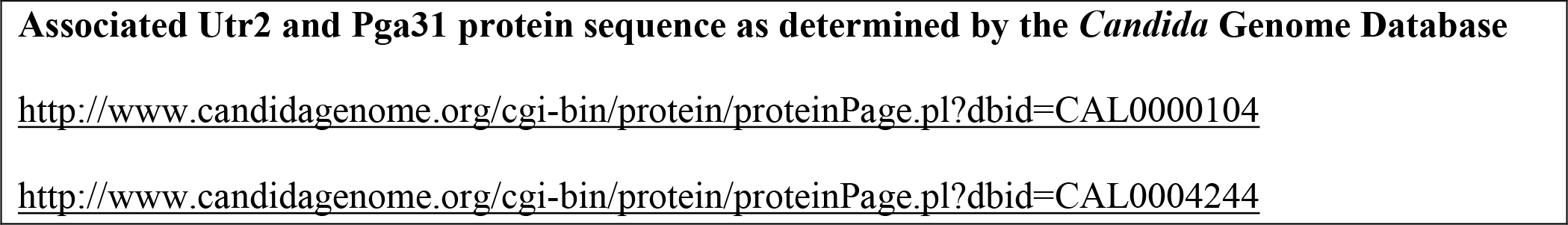
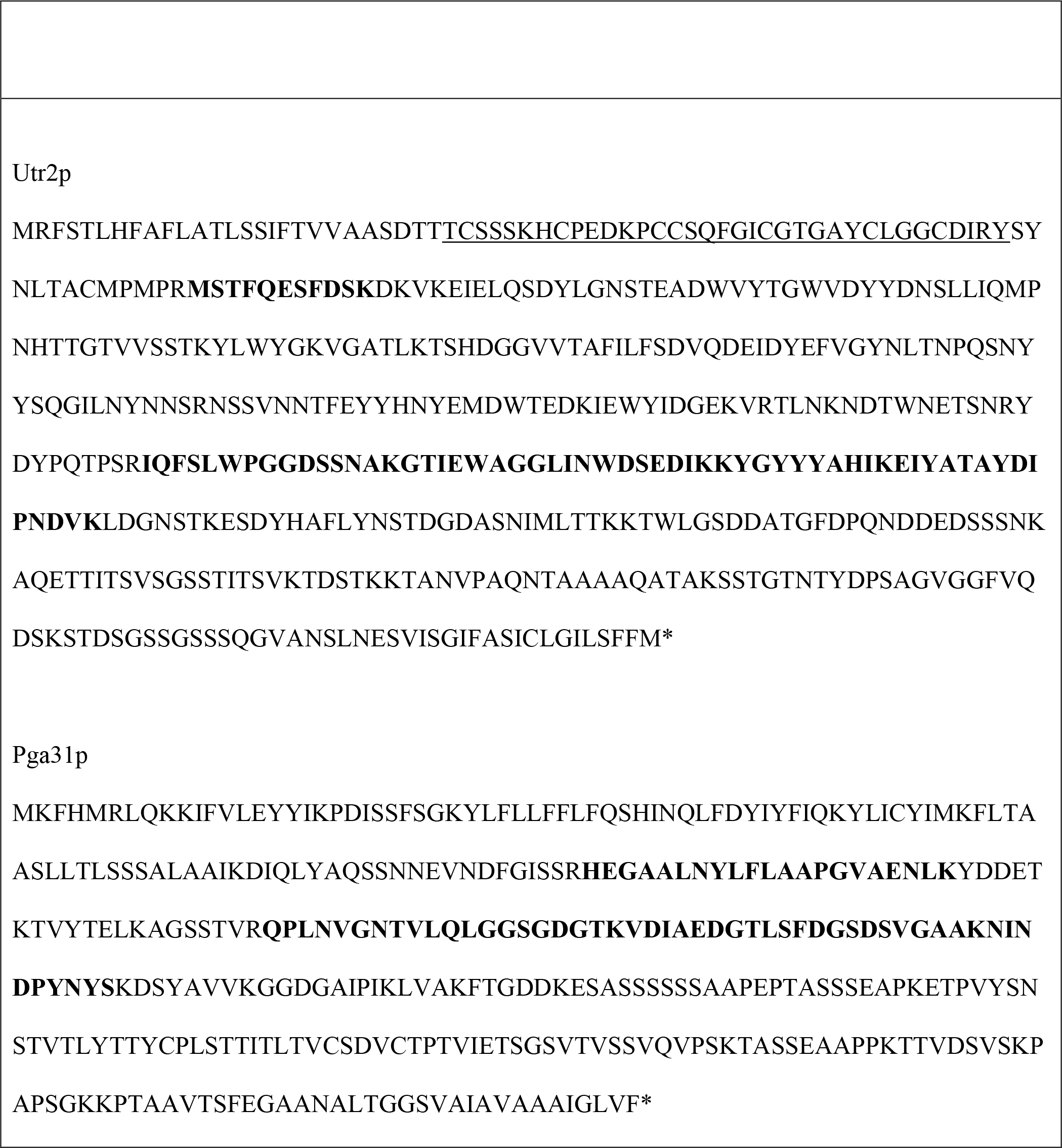
Mapping of the tryptic peptides identified in the cell wall proteome analysis onto C. albicans Utr2p and Pga31p sequences. Peptides are shown in bold within the amino acid sequences of each protein. The chitin binding region of Utr2, identified using SMART tool analysis software (http://smart.embl-heidelberg.de), is underlined.

### Isolation of *C. albicans* CWP specific recombinant antibody fragments from a human antibody library through biopanning

Phage display technology was employed to isolate Utr2 and Pga31 peptide specific antibody fragments from a human antibody library in single chain fragment variable (scFv) format. Following three rounds of selection using decreasing concentrations of biotinylated peptide antigen, two phage clones specific for the Pga31 peptide and 14 positive binders that recognised the Utr2 peptide were isolated and their unique scFv genes confirmed by DNA sequencing. These clones were reformatted into soluble single chain antibodies (scAbs) by cloning their respective VH-linker-VL region into the bacterial expression vector pIMS147 (30) to facilitate detection in biochemical assays and quantification of soluble expressed protein by *Escherichia coli* TG1 cells.

ScAbs from Pga31 biopanning reacted specifically to their peptide antigen and *C. albicans* wild type strain SC5314 hyphae (Fig. 1A & 1B) and bound total cell lysates of *C. albicans* SC3514 treated with or without 0.032 μg/ml caspofungin (Fig. 1C & 1D). An increase in scAb binding signal was recorded when lysate was prepared from the cells treated with caspofungin (Fig. 1C & 1D). This reinforced the hypothesis that Pga31 is overexpressed when the cells are grown in caspofungin, and that is involved in cell wall integrity (22). As a confirmatory control experiment, the scAbs did not bind to a *pga31*Δ mutant strain treated with/without caspofungin (Fig. 1E).

**Fig. 1.**
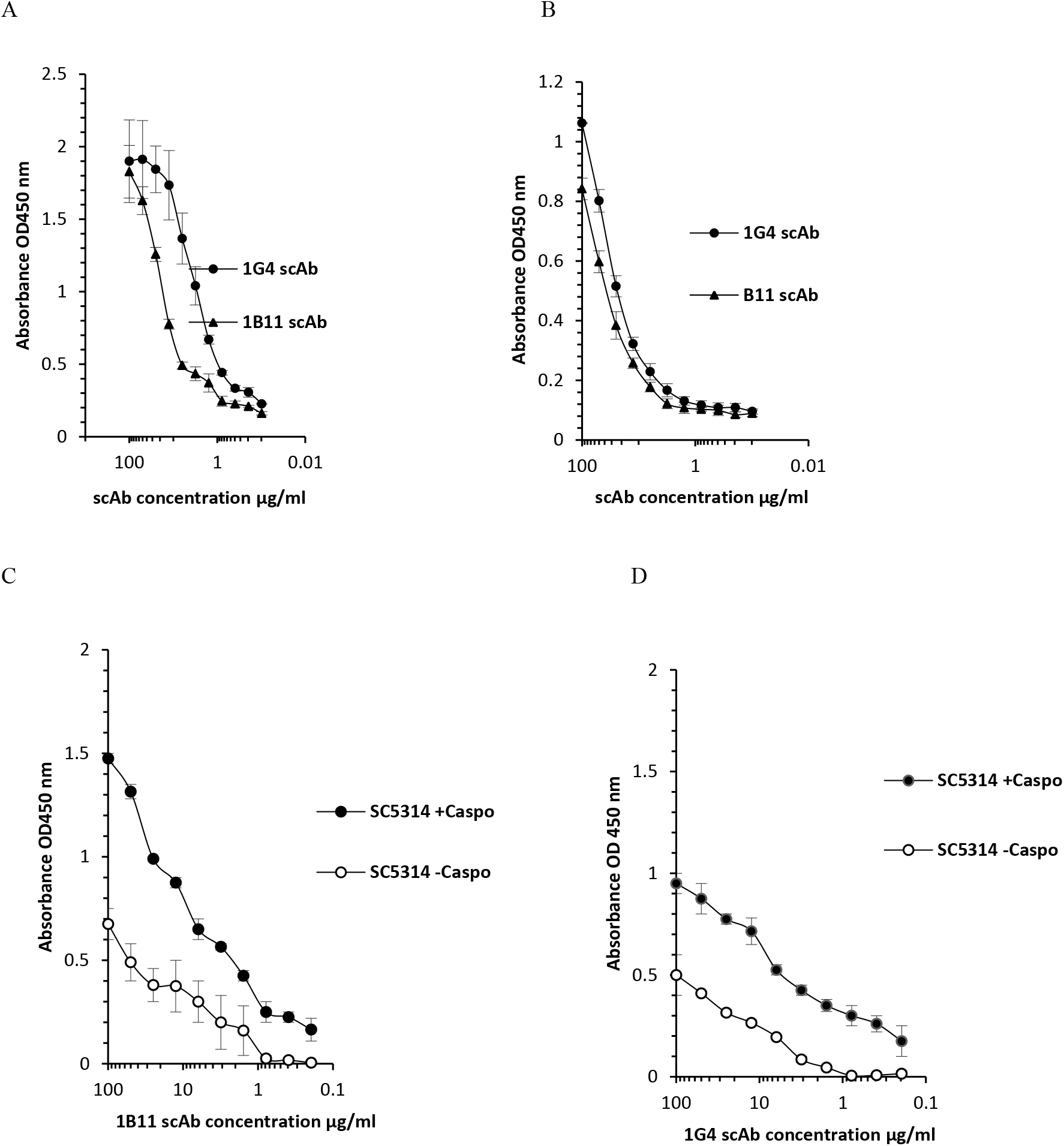
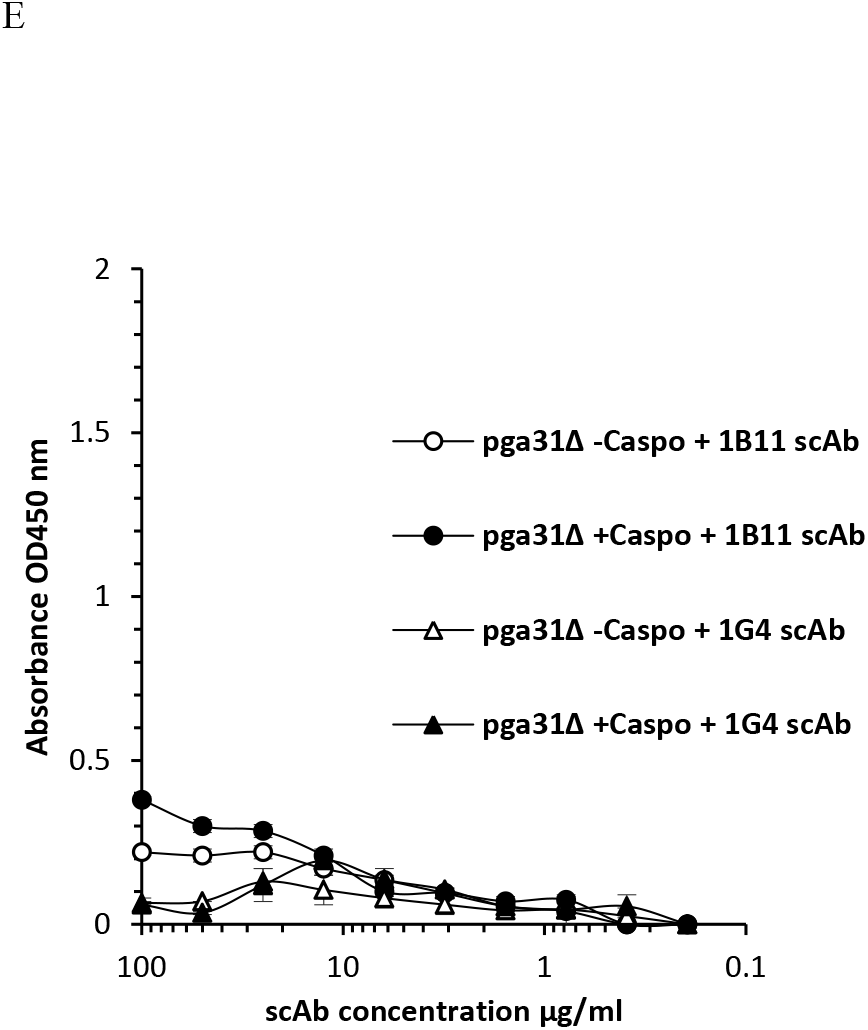
Pga31 scAbs binding profiles. (A) Antigen binding ELISA where wells are coated with Pga31 peptide–biotin conjugate. (B) scAb binding to C. albicans SC5314 hyphae (C) scAb 1B11 binding to total cell lysate of C. albicans SC5314 treated with or without 0.032 μg/ml caspofungin. (D) scAb 1G4 binding to total cell lysates of C. albicans SC5314 treated with or without 0.032 μg/ml caspofungin. (E) Lack of binding for 1B11 and 1G4 scAbs to the cell lysates of C. albicans pga31Δ mutant strain treated with or without 0.032 μg/ml caspofungin. Doubling dilutions of scAbs were added to the plates coated with Pga31 peptide conjugate, C. albicans SC5314 (-/+ caspofungin) or pga31Δ mutant (-/+ caspofungin) cell wall lysates and detected using an anti-human C kappa HRP conjugated secondary antibody. Values represent the mean absorbance OD450 nm readings (n=2, samples run in duplicate), error bars denote standard error of the mean (SEM).

The antigen binding of top four Utr2 specific scAbs 1B1, 1D2, 1F4 and 1H3, selected based on peptide-biotin conjugate ELISA signals, was confirmed (Fig. 2A) and these clones were tested for their ability to recognise Utr2 in cell lysate preparations of *C. albicans* SC5314 yeasts (Fig. 2B). A non-specific negative control scAb was unable to bind to *C. albicans* cell lysate (Fig. 2B). The surface exposure and epitope accessibility of the Utr2 antigen peptide was established, with all scAbs binding to *C. albicans* hyphal cells (Fig. 2C). When yeast cells were treated with 0.032 µg/ml caspofungin, an increase in scAb binding was observed when compared to fungal cells grown in the absence of the drug (Fig 2D-F). No scAb binding was seen when the *C. albicans* single mutant strain *utr2*Δ and triple mutant strain *utr2*Δ:*crh11*Δ:*crh12*Δ were used (Fig. 2G).

**Fig. 2.**
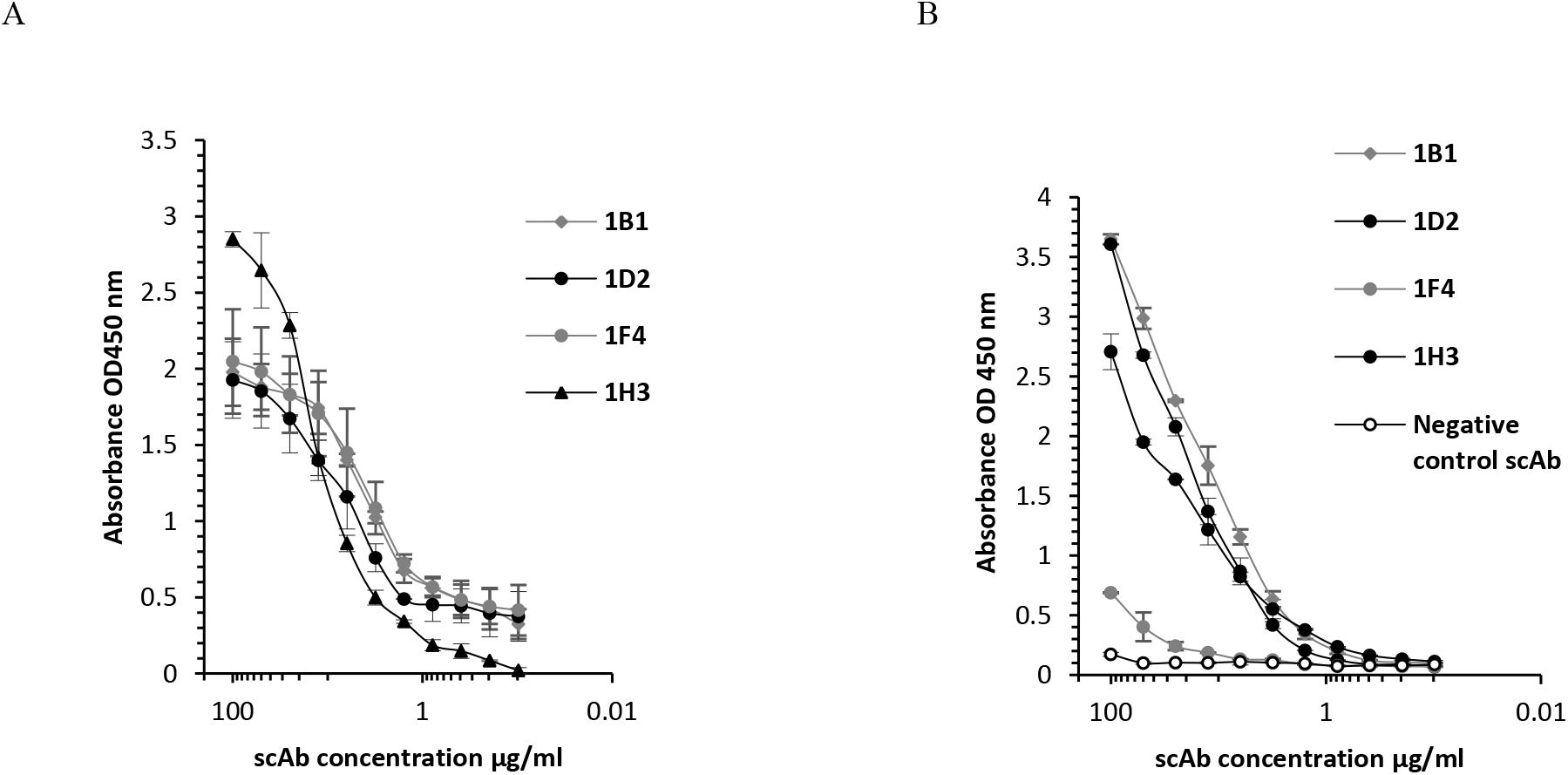
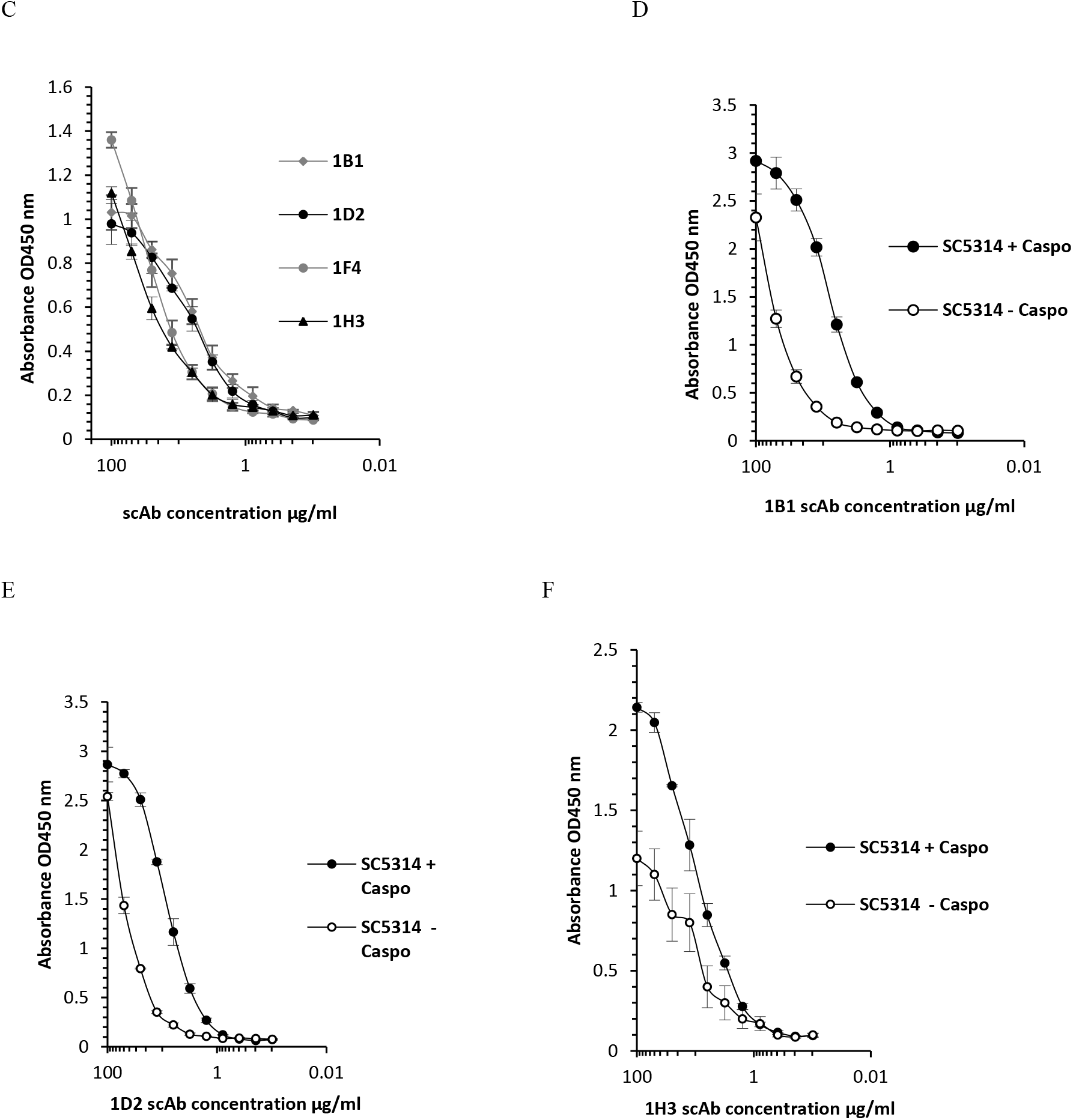
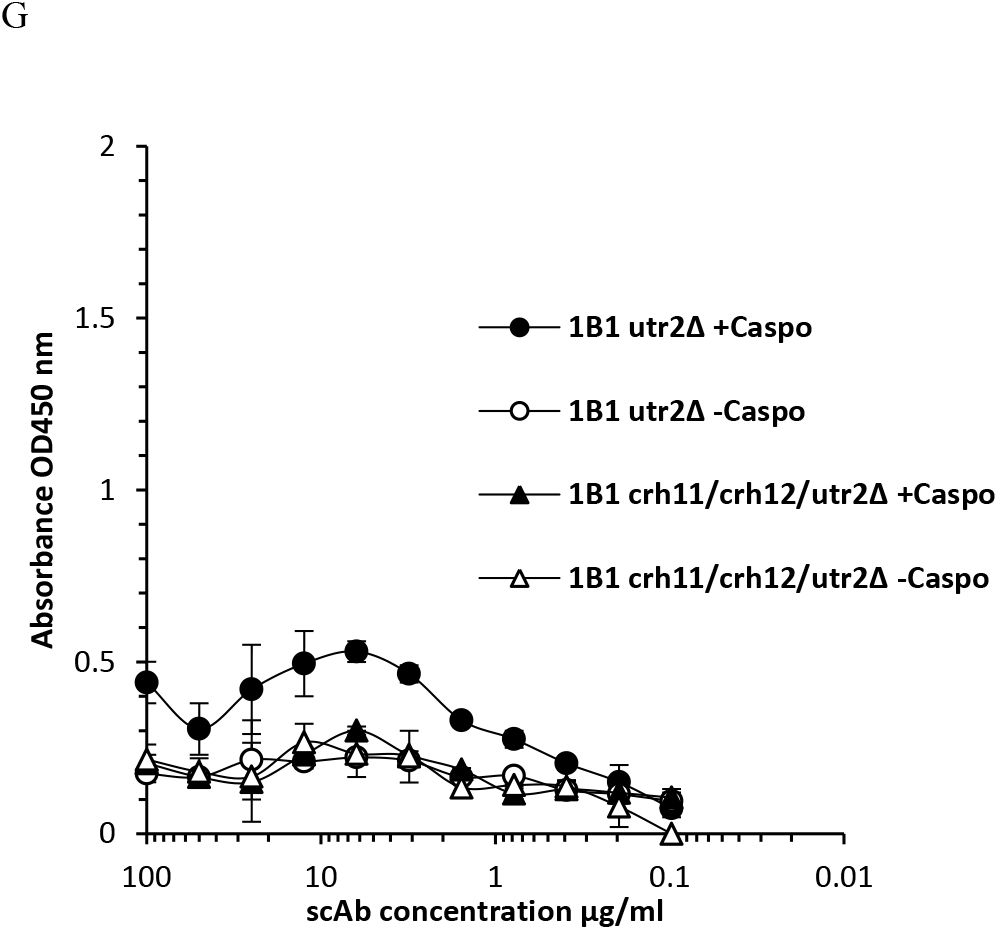
Utr2 scAb binding profiles. (A) Utr2 scAbs binding activity specific for the immobilised Utr2 peptide, (B) scAb binding using a cell lysate preparation of C. albicans SC5314 (C) scAb binding against C. albicans SC5314 hyphae (D-F) scAbs 1B1, 1D2 and 1H3 binding to C. albicans SC5314 yeast cells treated with or without 0.032 μg/ml caspofungin. (G) Representative scAb 1B1 binding to the cell lysates of a utr2 single mutant (utr2Δ) and triple mutant (utr2Δ/crh11Δ/crh12Δ) treated +/- caspofungin at 0.032 µg/ml. ScAbs 1H3, 1D2 and 1F4 also showed similar binding profiles with utr2Δ mutant strains, results not shown. Values represent mean absorbance OD450 nm readings (n=2, samples run in duplicate) Error bars denote standard error of the mean (SEM).

### Reformatting scAbs into human-mouse chimeric IgGs for *in vitro* and *in vivo* validation studies

Utr2 scAbs 1D2, 1H3 and Pga31 scAb 1B11 were selected for IgG reformatting based on their binding interactions with target peptides and *C. albicans* cells, and protein expression levels. The VH and VL domain genes of the Utr2 scAbs 1D2, 1H3 and Pga31 scAb 1B11 were cloned into a dual plasmid eukaryotic expression system encoding mouse IgG2a and kappa constant domain genes and the resultant recombinant chimeric mAbs were expressed transiently in HEK293-F cells. The presence of functional, protein A affinity-purified mAbs 1B11 and 1D2, were confirmed by antigen binding ELISA with no cross-reactivity observed to unrelated peptide sequences (Fig. 3A & 3B). MAb 1H3, which was selected against the Utr2 peptide, appeared to also recognise a peptide sequence selected as a surface exposed region from the *C. albicans* cell wall protein Phr2 (Fig. 3C). The reason for this is unclear.

**Fig. 3.**
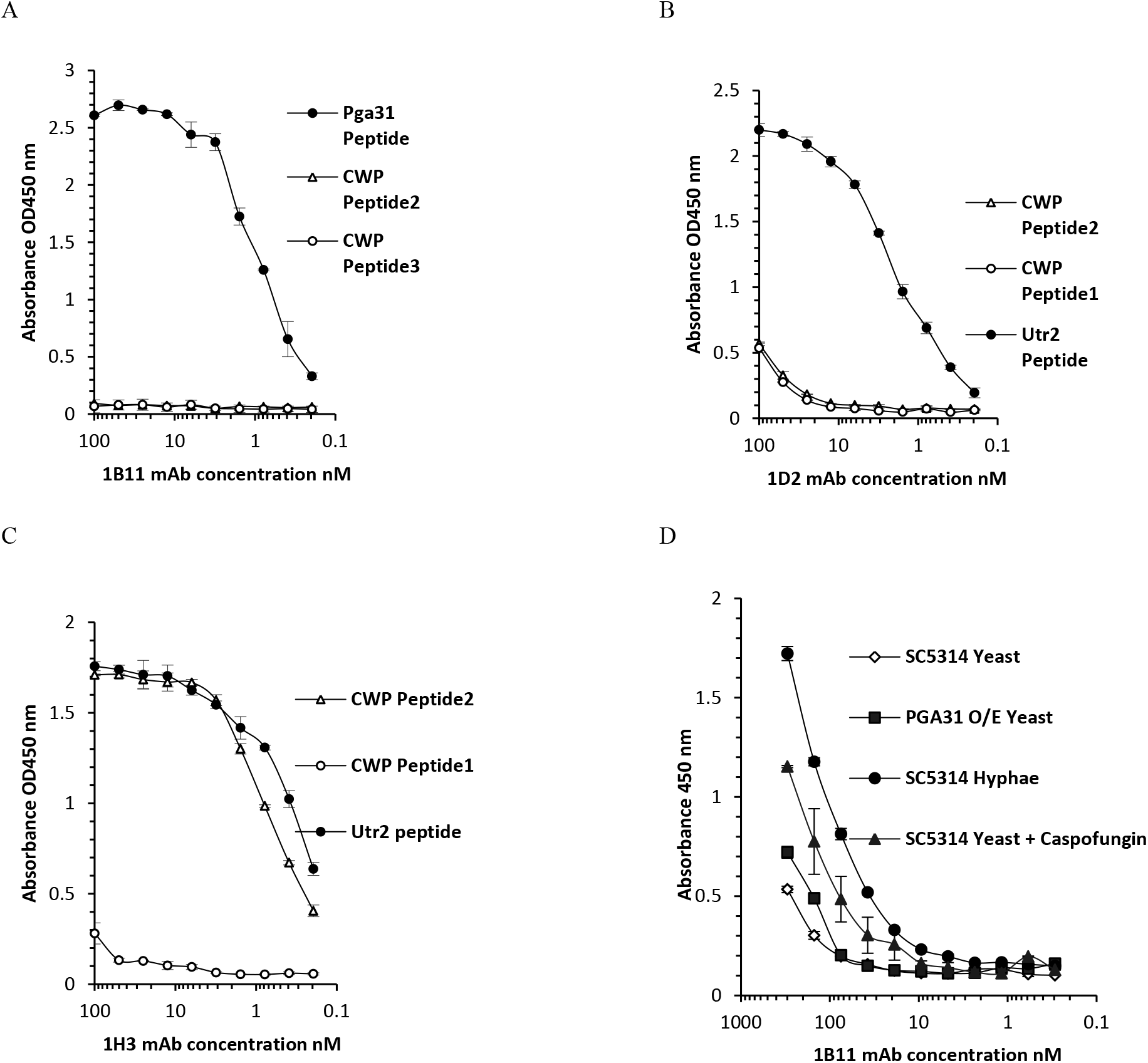
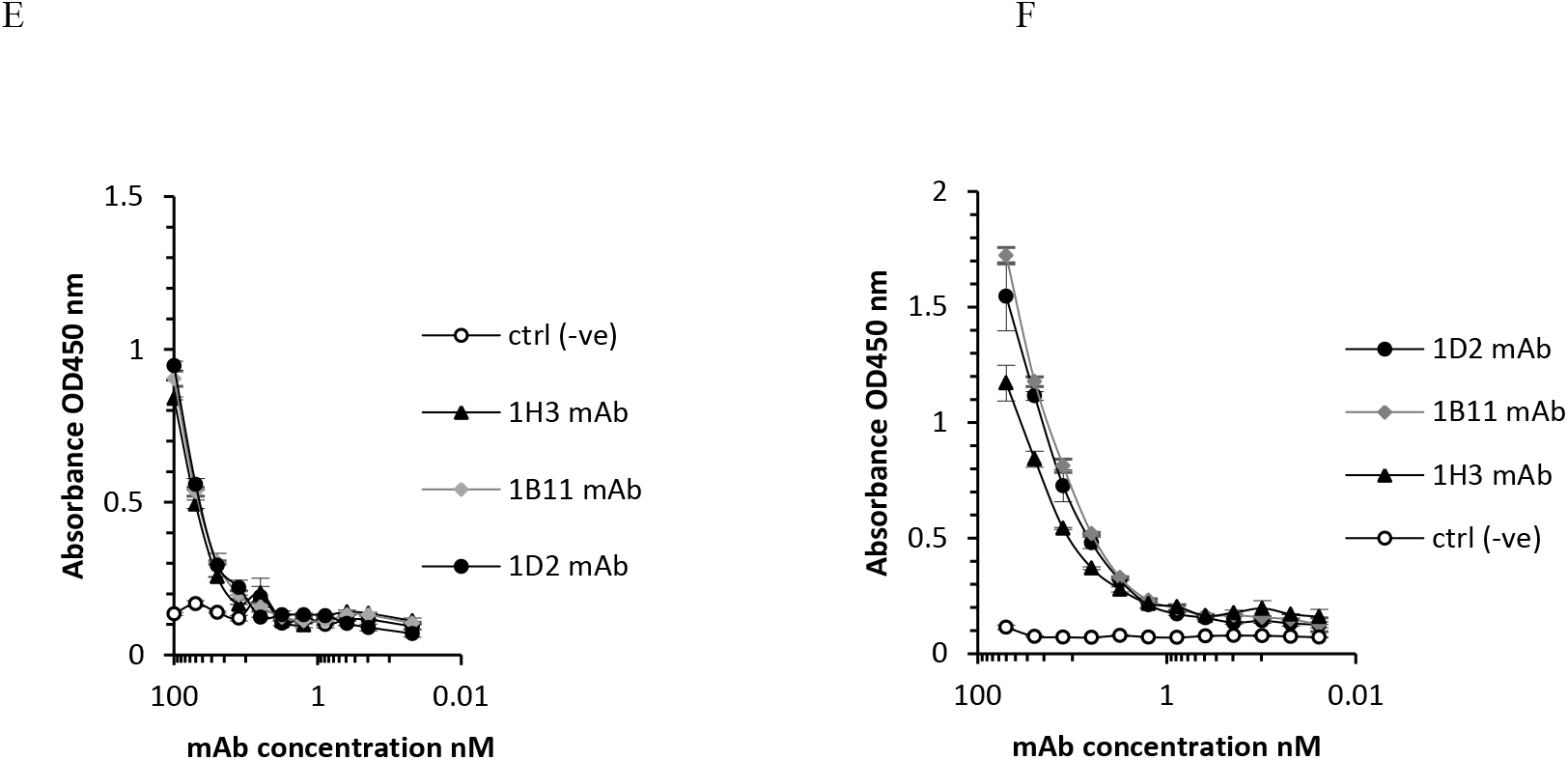
Reformatted Pga31 and Utr2 mAb binding profiles. (A) 1B11 mAb binding to streptavidin captured biotinylated Pga31 peptide whilst assessing also the cross-reactivity for other C. albicans cell wall proteins (CWP2 and CWP3) using peptide antigens isolated following proteome analysis (B,C) 1D2 and 1H3 mAbs binding to streptavidin captured biotinylated Utr2 peptide and assessment of the cross-reactivity for other C. albicans cell wall protein peptide antigens as above (D) 1B11 mAb binding to C. albicans SC5314 yeasts, a *PGA31* over-expressing strain, *C. albicans* SC5314 hyphae, and C. albicans SC5314 yeasts treated with 0.032 µg/ml caspofungin (E) 1B11, 1H3 and 1D2 mAbs binding to *C. albicans* SC5314 yeast (F) 1B11, 1H3 and 1D2 mAbs binding to C. albicans SC5314 hyphae. The binding of mAbs to C. albicans peptides and whole cells was detected and quantified using an anti-mouse IgG Fc region specific HRP conjugated secondary antibody and the values plotted represent mean absorbance readings at OD450 nm (n=2, samples run in duplicate). Error bars indicate standard error of the mean (SEM)

The conversion of scAbs into a bivalent IgG format significantly increased (possibly in part *via* avidity) the relative binding affinities of all three lead antibodies. Their EC_50_ values (antibody concentration required to achieve 50% reduction in maximum absorbance) were calculated by extrapolating values obtained from direct antigen binding plots (Fig. 1A, 2A and 3A-C). The calculated EC_50_ for 1H3 scAb from the peptide binding assay was 175 nM, whereas the reformatted 1H3 mAb achieved half maximal binding at 400 pM, an apparent 400-fold improvement in functional affinity. Similarly, the EC_50_ values obtained for 1D2 scAb and mAb were 80 nM and 2 nM respectively. For the Pga31 clone 1B11, mAb reformatting resulted in 600-fold improvement in antigen binding compared to the parental scAb clone, with estimated EC_50_ values of 600 pM and 375 nM respectively.

In a whole cell binding ELISA, Pga31 mAb 1B11 preferentially recognise wild type *C. albicans* (SC5314) hyphae compared to the yeast form (Fig. 3D), suggesting a morphology dependent binding function which could signify increased epitope accessibility in this phenotype. Although relatively poor binding activity was observed for the yeast form, when the cells were stressed with caspofungin treatment, an enhancement in mAb targeting was seen (Fig. 3D). 1H3 and 1D2 mAbs were able to bind SC5314 cells immobilised on maxisorbant plates (Fig. 3E and 3F), however similar to Pga31 mAb 1B11, Utr2 mAbs were also seen to display increased binding activity for the hyphae when compared to the yeast cells.

### Immunofluorescence staining of *C. albicans* using Utr2 and Pga31 antibodies

Immunofluorescent microscopy using cell wall protein specific antibodies 1D2, 1H3 and 1B11 demonstrated specific and distinct binding patterns on *C. albicans* cells. Antibodies 1B11 (anti-Pga31) and 1D2 (anti-Utr2) bound to *C. albicans* SC5314 hyphae but exhibited little or no binding to mother yeast cells (Fig. 4). 1D2 staining was visible across the entire hyphal surface, whereas 1B11 showed more punctate binding in distinct hyphal regions (Fig. 4A). The cross-reactive 1H3 mAb (anti-Utr2), appeared to bind specifically to the apical tip of growing hyphae (Fig 4A). When *C. albicans* SC5314 yeast cells were stained using anti-Utr2 and anti-Pga31 mAbs, a punctate binding pattern was observed on the surface of a limited number of cells (Fig. 4B). In contrast, when yeast cells were pre-treated with 0.032 µg/ml caspofungin, strong binding, again punctate in nature, was seen in a large proportion of cells at multiple sites including regions of possible bud emergence (Fig 4C). Similar to hyphal staining, 1D2 mAb produced a distinct binding pattern in budding yeasts, where staining was localised to the emerging daughter cells in areas where new cell wall is being produced (Fig. 4C). The negative control mAb did not show any staining of caspofungin treated or untreated cells (Fig. 4B). In summary, all three antibodies displayed a morphology-specific binding pattern, including increased binding to yeast cells treated with caspofungin, supporting the earlier ELISA data (Fig. 3).

**Fig. 4.**
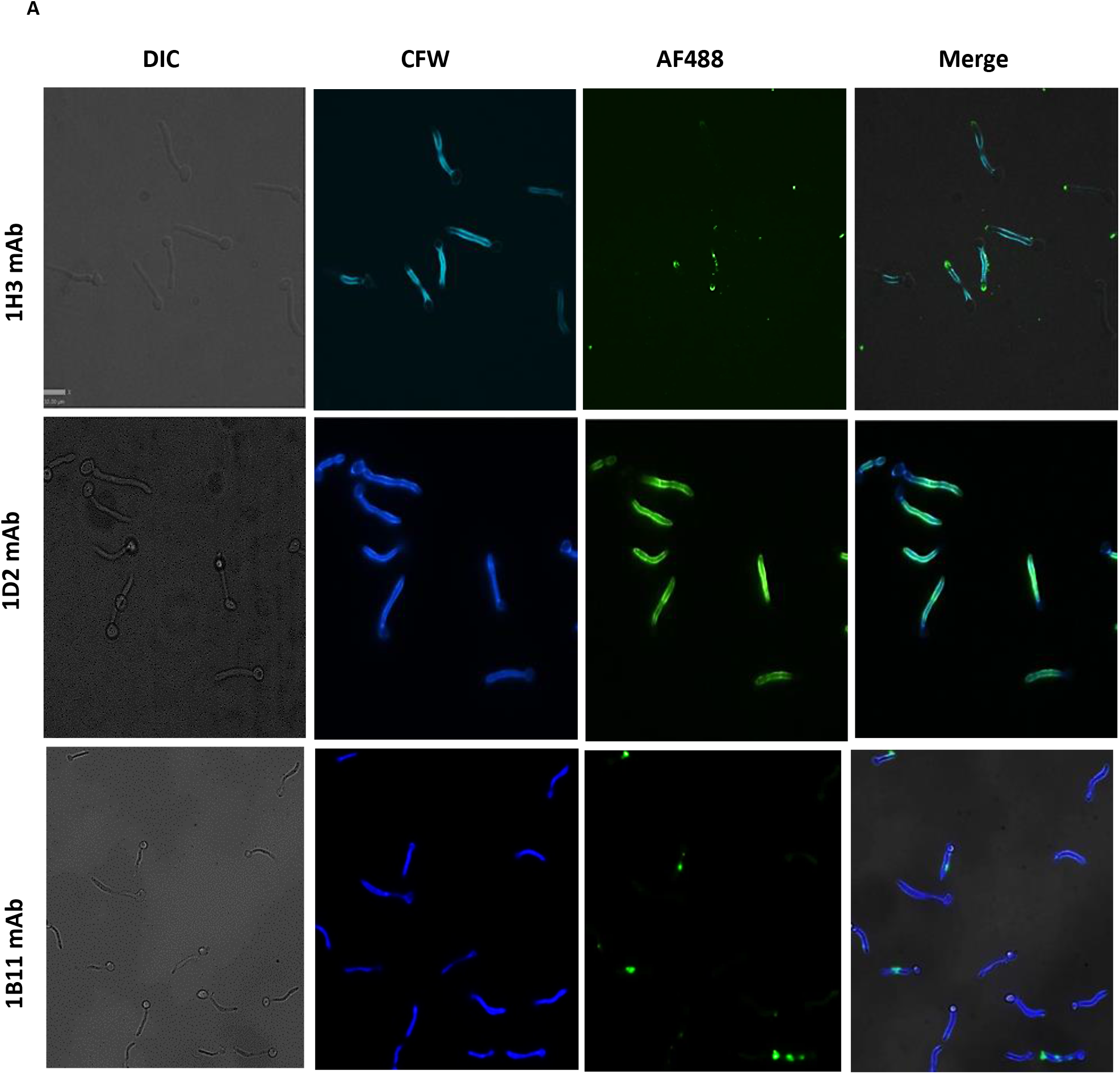
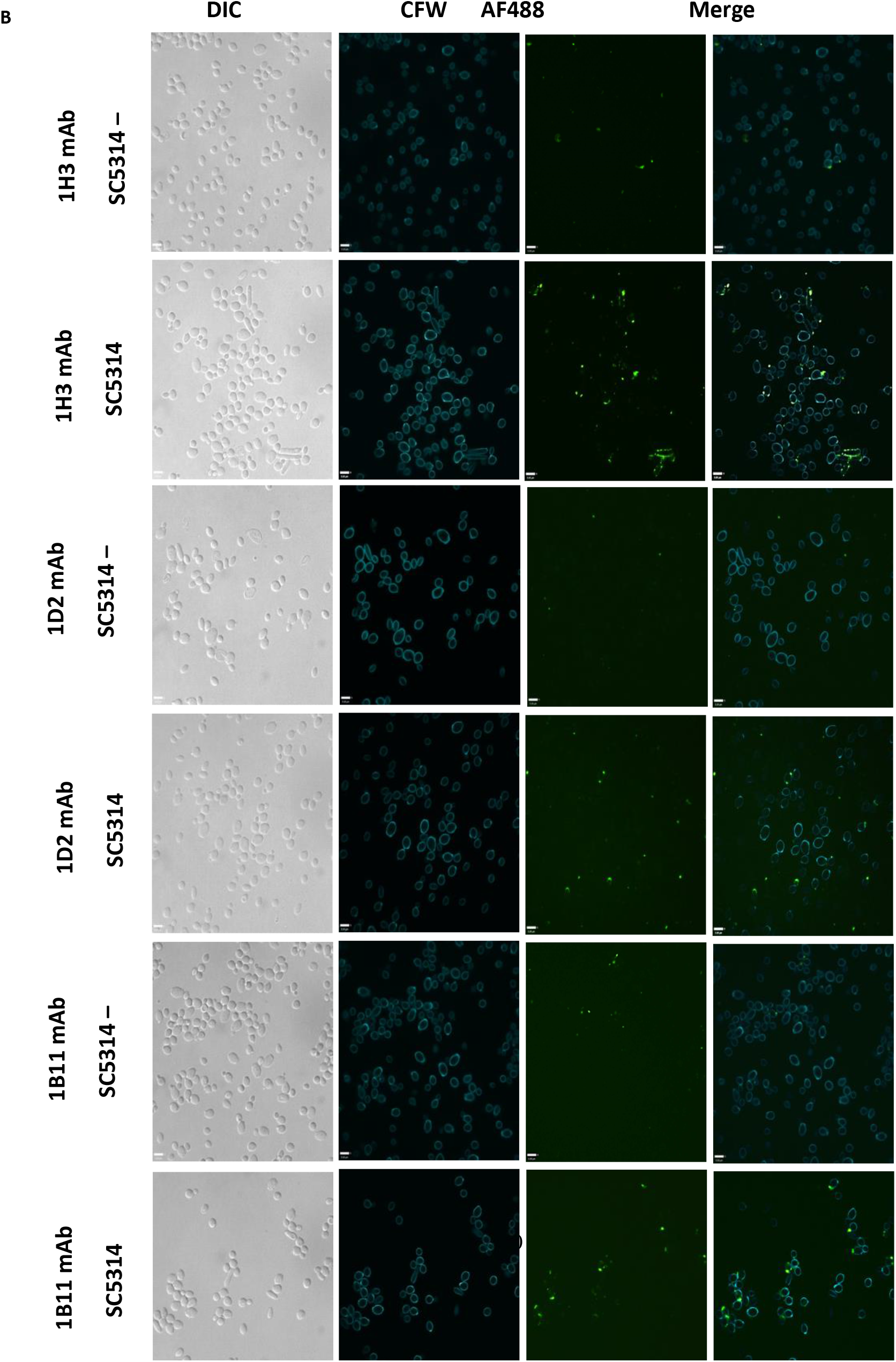
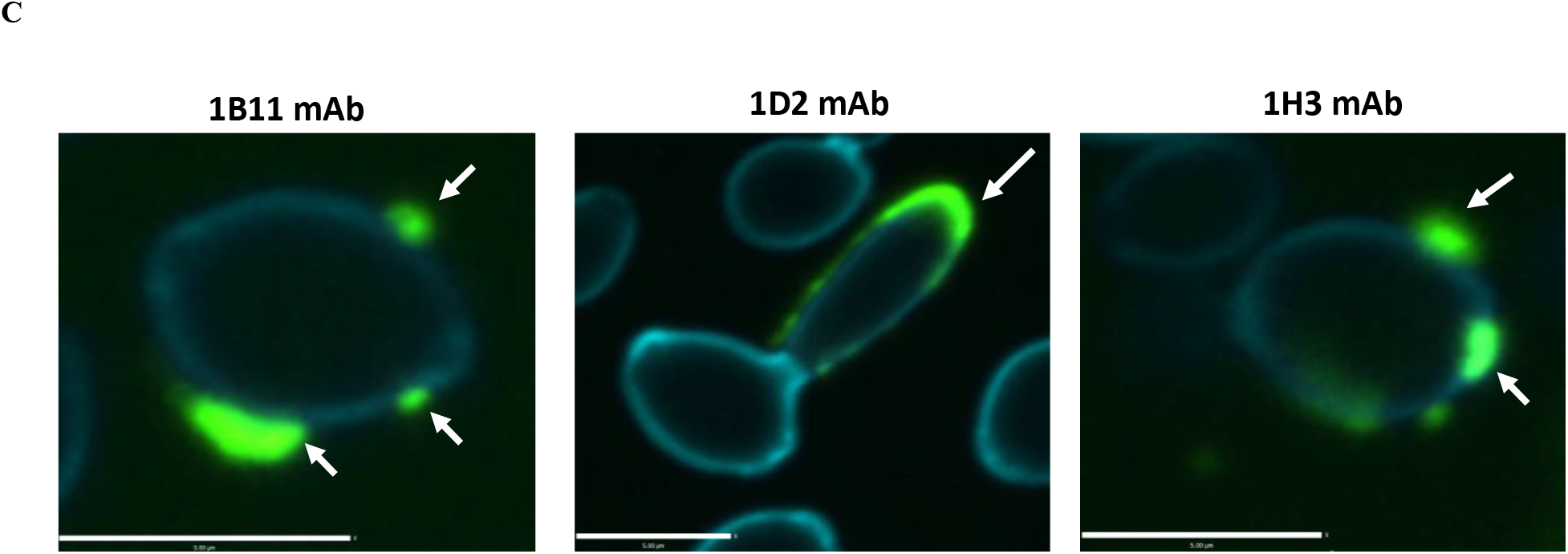
Utr2 and Pga31 antibody binding to C. albicans yeast and hyphae. (A) C. albicans SC5314 hyphae immunostaining with 1H3, 1D2 or 1B11 antibodies. Calcofluor white (CFW) was used to stain cell wall chitin and Alexa Fluor 488 (AF488) conjugated goat anti-mouse IgG2a antibody was used to detect CWP-specific mAb binding. Green fluorescence indicates antibody binding on the cell surface and distinct binding patterns were observed with the three test mAbs. The anti-Utr2 antibody 1H3 binds at the apical tip of growing hyphae. In contrast, the second Utr2 mAb 1D2 displayed uniform binding along the hypha. The anti-Pga31 mAb 1B11 had a more localised binding pattern with binding to a single major location on the growing hypha. *(B)* C. albicans *SC5314 yeast cells treated with/without caspofungin (0.032 µg/ml) and immunostained with 1H3, 1D2 or 1B11 antibodies or a negative control mouse Ig2a antibody. Increased mAb binding was observed in caspofungin treated cells, mostly as a punctate binding pattern around the poles of buds*. *(C) The anti-Utr2 mAb 1D2 displays distinct binding, with intense staining localised around zones of polarised growth and away from the mother cell. The second anti-Utr2 mAb 1H3 and anti-Pga31 mAb 1B11 showed punctate binding on the cell surface (indicated by white arrows)*.

### Macrophage Interaction assay

To evaluate the ability of CWP-specific mAbs to potentially confer protection in an infection model, a macrophage interaction assay was performed using anti-Pga31 or anti-Utr2 mAbs as opsonising agents for immune cell recruitment and mediation of phagocytosis. *C. albicans* SC5314 untreated or pre-coated with test mAbs or a commercially sourced anti-*Candida* IgG were used to challenge mouse J774.1 macrophage-like cells. The outcomes were visualised by live cell video microscopy. Engulfment time (time taken for the macrophages to engulf *C. albicans* cells) and the length of intracellular hyphae were determined.

Cells pre-incubated with mAbs and mouse IgG control were engulfed significantly more rapidly when compared to *C. albicans* without antibody pre-treatment (Table 2). The vast majority (95%) of fungal cells were engulfed within 7 min compared to 10 min for untreated cells (Supplementary Fig. S1A-D). When incubated with anti-Pga31 or anti-Utr2 mAbs all fungal cells were engulfed by 12 min; however, in the case of untreated *C. albicans*, this took 15 min (Supplementary Fig S1A). These data suggest that anti-CWP mAbs influence macrophage behaviour by targeting fungal cells for opsonophagocytosis.

**Table 2.** Time taken for J774.1 mouse macrophages to engulf anti-Candida mAb-treated or untreated C. albicans SC5314. At least twenty-five macrophages were selected at random per video to determine the engulfment time. Data represents average time taken ± SD (min) The length of intracellular hyphae at multiple incubation times was also analysed from microscopy videos (Fig. 5). Measurements were taken from the neck of the hypha to the apical tip at 60 & 90 min following co-incubation with macrophages. Our data shows that intracellular hyphae at 60 min were significantly shorter for *C. albicans* cells pre-incubated with anti-Pga31 mAb compared to all other treatment groups (P = < 0.0001) (Fig 5A). *C. albicans* cells pre-treated with the positive control IgG were also shorter compared to untreated, but the difference was only significant at 90 min (Fig. 5A & 5B).

| Treatment group | Average engulfment time<br>mean ±SD (min) |
| --- | --- |
| SC5314 (wildtype) | 5.64 ± 2.59 |
| SC5314+anti-Pga31 mAb | 4.18 ± 1.80 |
| SC5314+anti-Utr2 mAb | 4.28 ± 1.96 |
| SC5314+IgG positive control mAb | 4.29 ± 2.43 |

**Fig. 5.**
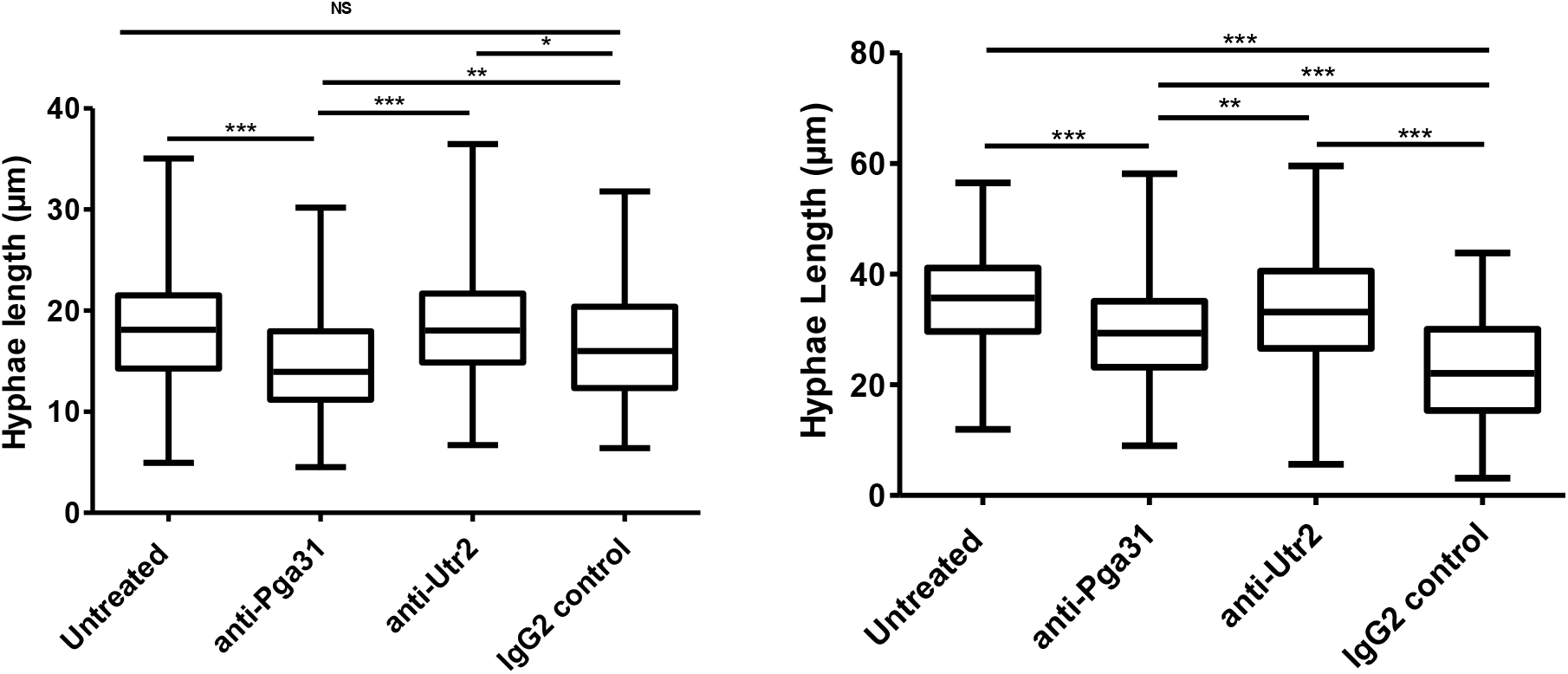
Length of intracellular hyphae at 60 and 90 min. Intracellular hyphal lengths were measured (µm) following C. albicans uptake by J774.1 mouse macrophage at 60 (A) and 90 (B) min. Twenty-five macrophages were selected at random per video to measure the length of intracellular hyphae at two time points. Statistical significance was determined by Kruskal-Wallis test with Dunn’s multiple comparison test *P < 0.05, **P < 0.01, ***P < 0.005.

### Testing the therapeutic efficacy of anti-Utr2 and anti-Pga31 mAbs in a *C. albicans* mouse infection model

A series of *in vivo* mouse infection studies were conducted to evaluate the protective effect of Pga31 and Utr2 mAbs in a disseminated candidiasis model, with the efficacy measured by determining the organ fungal burden and mouse survival. In prophylactic study 1, mice were pre-treated with 15 mg/kg of the test mAbs (including an isotype control), 3 h prior to IV administration of *C. albicans* SC5314, followed by a second dose of mAb 24 h post-infection. All treatments were administered intraperitoneally (IP) in 150 μl saline. In the anti-Pga31 mAb treated group, 67% of mice survived four days post-infection, whereas the Utr2 mAb conferred 33% protection (Fig. 6A). Saline only and isotype control groups did not show any survival benefit at 4 days. Comparing across all groups there were significant differences in survival (*P*=0.045, Kaplan-Meier log-rank statistics).

**Fig. 6(A).**
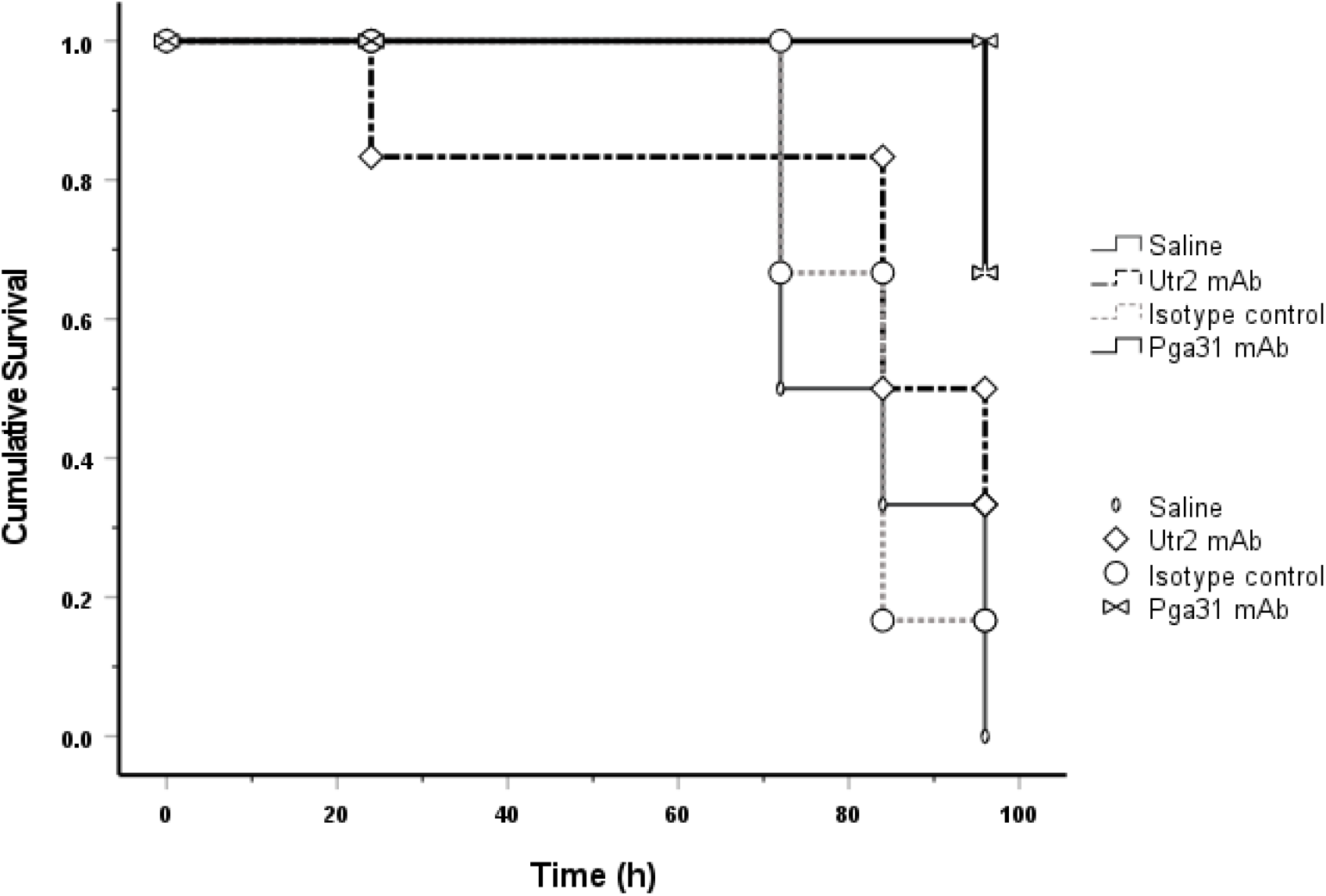
Kaplan-Meier survival curve representing the treatment effect of test mAbs and control groups over 96 h in Study 1. Pga31 mAb 1B11 (15 mg/kg), Utr2 mAb 1D2 (15 mg/kg), isotype control IgG (15 mg/kg) or saline were administered IP to mice 3 h pre and 24 h post-infection with C. albicans SC5314. Mice (n=6/group) surviving the study period were treated as censored data for analysis and statistical significance of survival between groups was determined using the log-rank test.

Comparing the percentage weight change (day 0-2, supplementary Table S1) and kidney fungal burdens (Fig. 6B), the differences were again statistically significant between some groups (*P* <0.001 Kruskal-Wallis Dunn’s multiple comparison test). In particular, when comparing the fungal kidney burden of the saline only group to Pga31 mAb treated group (*P*=0.004) and weight change (*P*=0.011). Utr2 treated group showed a significant difference for weight change only (*P*=0.023). There was no difference in kidney fungal burdens between the isotype control IgG-treated mice and saline-treated mice (*P*>0.999).

**Fig. 6(B).**
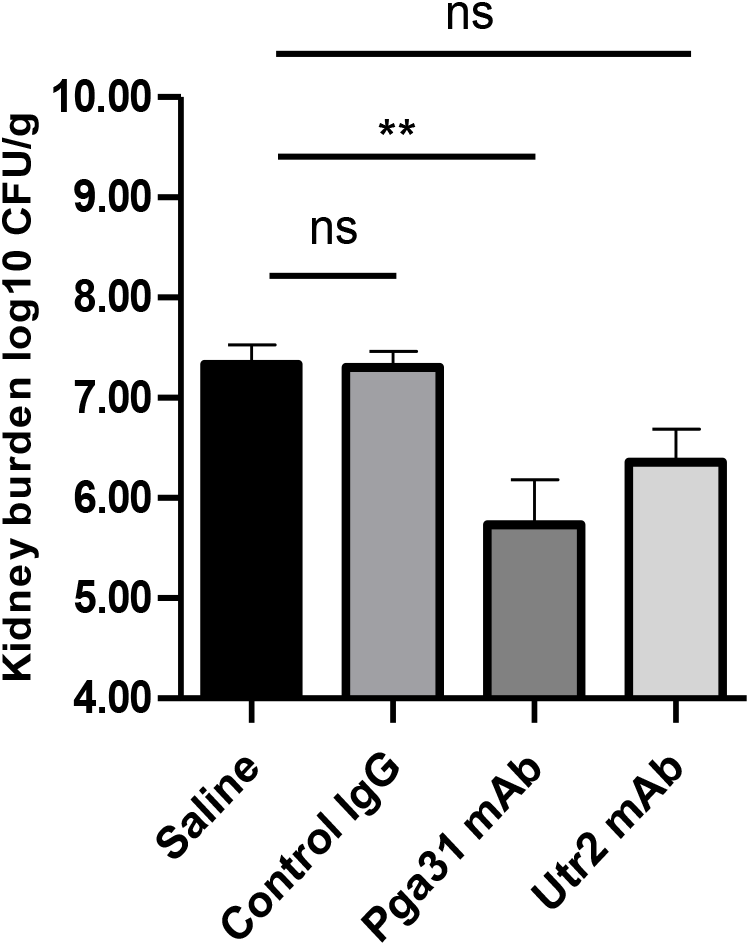
Mean fungal burdens in kidneys on day 4 post-infection in Study 1. Isotype control IgG (15 mg/kg IP), Pga31 mAb 1B11 (15 mg/kg IP) and Utr2 mAb 1D2 (15 mg/kg IP) were administered 3 h pre- and 24 h post-infection. Error bars denote standard deviation. A significant difference was observed for kidney burden between the saline treatment group and the 1B11 mAb therapy group (n= 6 mice/group; P=0.004, Kruskal-Wallis Dunn’s multiple comparison test)

A second study was conducted to test the effectiveness of multiple dosing of antibodies. A single 12.5 mg/kg dose was administered prophylactically followed by two treatment doses at 24 and 72 h post infection. All treatments were administered intraperitoneally (IP) in 150 μl saline and the survival of mice treated with Pga31 and Utr2 mAbs were compared with that of an isotype control. Mice were monitored and weighed every day and were culled on day 6 post-infection and fungal burdens in several organs including the kidneys, brain and spleen were determined as before. While mouse survival of 83% was achieved with the Pga31 mAb following a second dose 72 h post infection (vs 66.7% for study 1), no further improvement was observed for the Utr2 mAb (33 % survival in study 1 and 2) (Fig. 7). Isotype control IgG showed no therapeutic effect and the differences between various groups are statistically significant (*P* <0.001, log-rank test). A one log drop in kidney fungal burden, representing killing of fungi or inhibition of cell division, for the Pga31 mAb treated group was achieved when compared to the isotype control group (Pga31 mAb = 5.5. log_10_ CFU/g vs isotype control = 6.8 log_10_ CFU/g); however, there was little or no difference in mean fungal counts for the Utr2 mAb group (Utr2 mAb = 6.7 log_10_ CFU/g vs isotype control = 6.8 log_10_ CFU/g) (Table 3). Similarly, fungal burden in associated organs including the brain and spleen was also reduced in test antibody groups, with Pga31 mAb showing an improved therapeutic effect when compared with the Utr2 mAb.

**Fig. 7.**
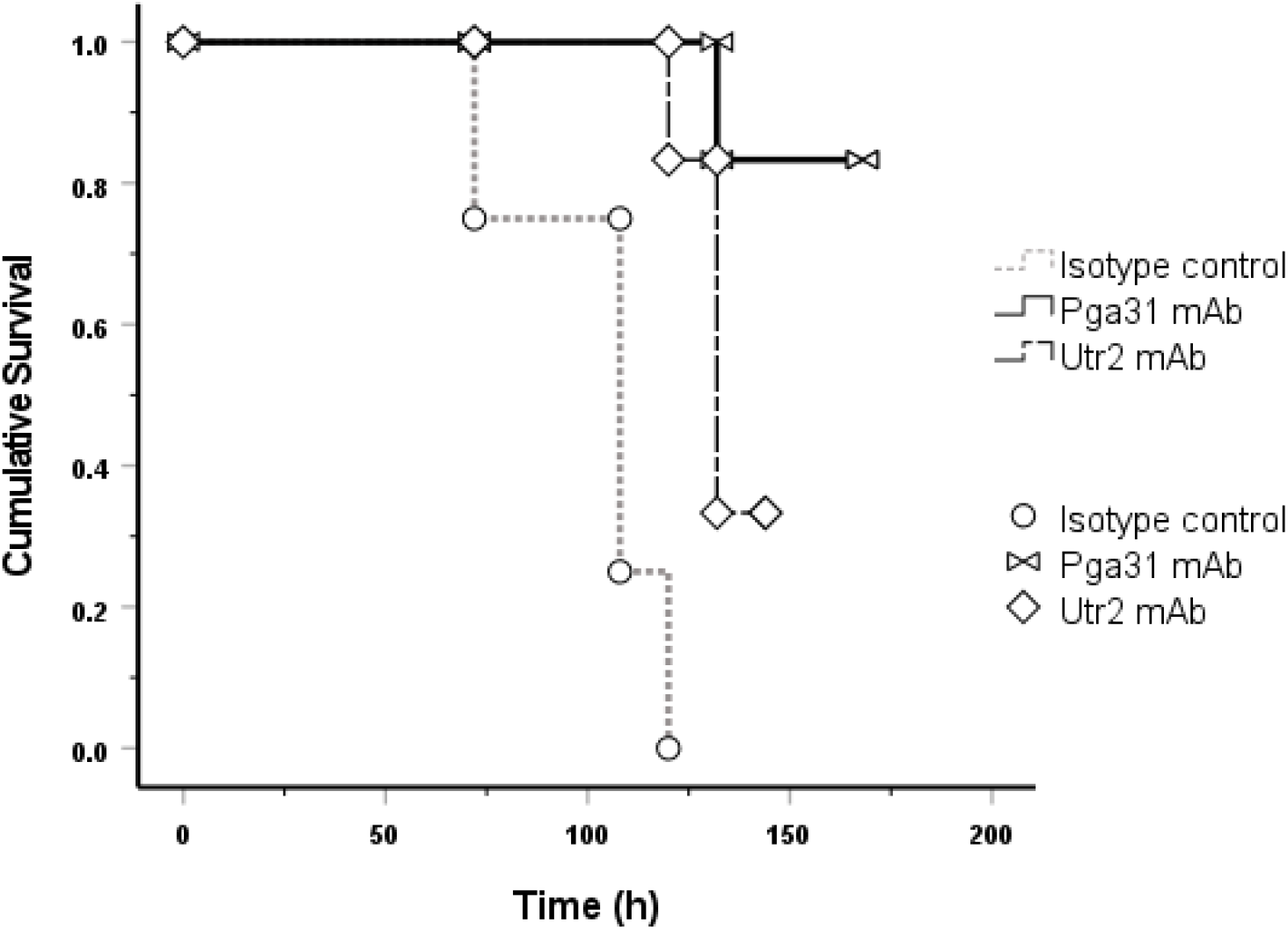
Kaplan-Meier survival curve representing the treatment effect of test mAbs and control groups 6 days post-infection (Study 2). Pga31 mAb 1B11 (12.5 mg/kg), Utr2 mAb 1D2 (12.5 mg/kg) or isotype control IgG (12.5 mg/kg) were administered IP in mice 3 h pre and 24 h and 48 h post-infection with C. albicans SC5314. The difference between the three groups (n= 6 mice/group) is statistically highly significant (P<0.001, Kaplan-Meier log-rank test)

**Table 3.**
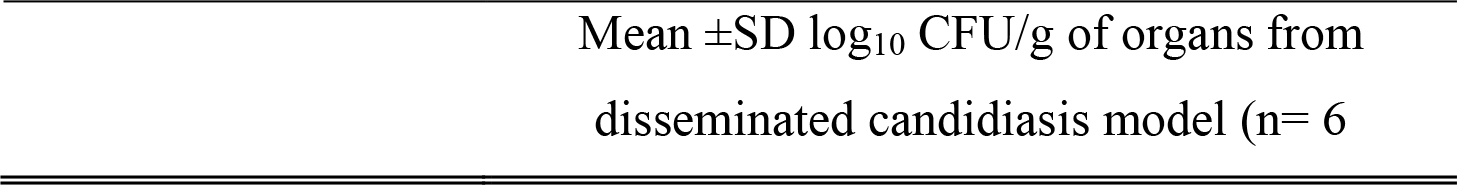
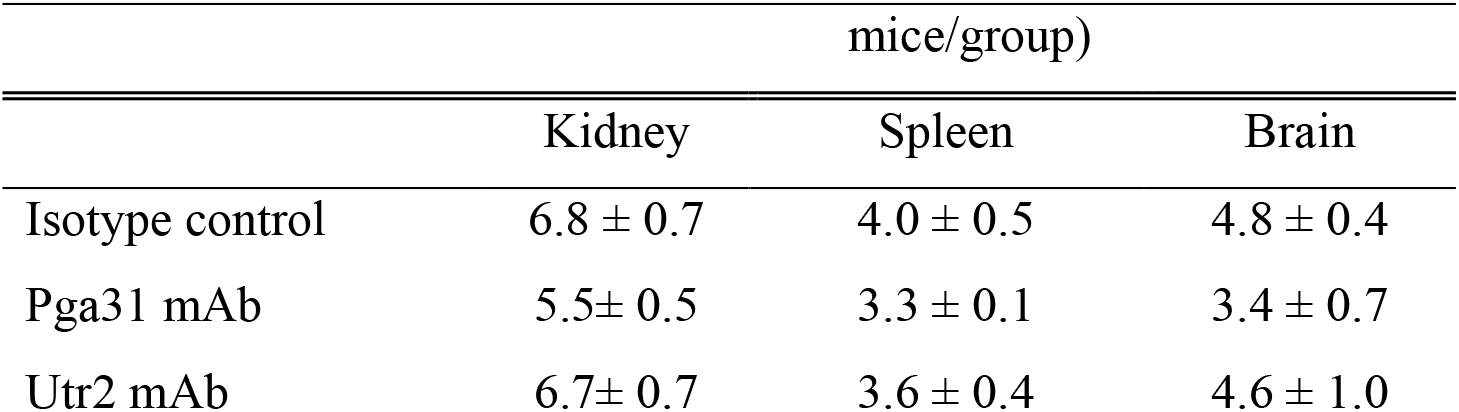
Mean fungal burdens in mouse organs at 6 days post-infection (Study 2). Control IgG (12.5 mg/kg), Pga31 mAb 1B11 (12.5 mg/kg IP) and Utr2 mAb 1D2 (12.5 mg/kg IP) were administered IP 3 h pre-infection and 24 h and 72 h post infection (n= 6 mice/group). Statistical significance was achieved for fungal counts in the kidneys only (P=0.02, Kruskal-Wallis Dunn’s multiple comparison test).

Finally, to compare the protective effect of test antibodies as a prophylactic agent vs treatment, a follow-up study (Study 3) was conducted. In the prophylactic arm a single dose of each test antibody was administered 3 h before infection followed by two doses at 24 h and 72 h post infection. For the treatment only arm, mAbs were given 24 h and 72 h post infection and the fungal burdens in the kidneys of various groups were compared with groups of mice receiving saline or caspofungin (1 mg/kg) (Fig. 8). The Pga31 mAb prophylactic arm significantly reduced the fungal burden in the kidneys of animals, 7 days post infection, which was similar to the levels achieved with caspofungin treatment (Pga31 mAb = 2.22 log_10_ CFU/g and caspofungin = 1.98 log_10_ CFU/g vs saline = 4.46 log_10_ CFU/g, *P*=0.002, Kruskal-Wallis test). Mice receiving only two doses of Pga31 mAb post-infection, also had reduced fungal burden in their kidneys (3.22 log_10_ CFU/g). Interestingly, for the Utr2 mAb groups, the treatment arm where the test antibody was administered 24 h and 72 h post infection had reduced burden in the kidneys as opposed to the prophylactic arm (Utr2 mAb treatment arm = 2.47 log_10_ CFU/g vs prophylactic arm = 3.16 log_10_ CFU/g).

**Fig. 8.**
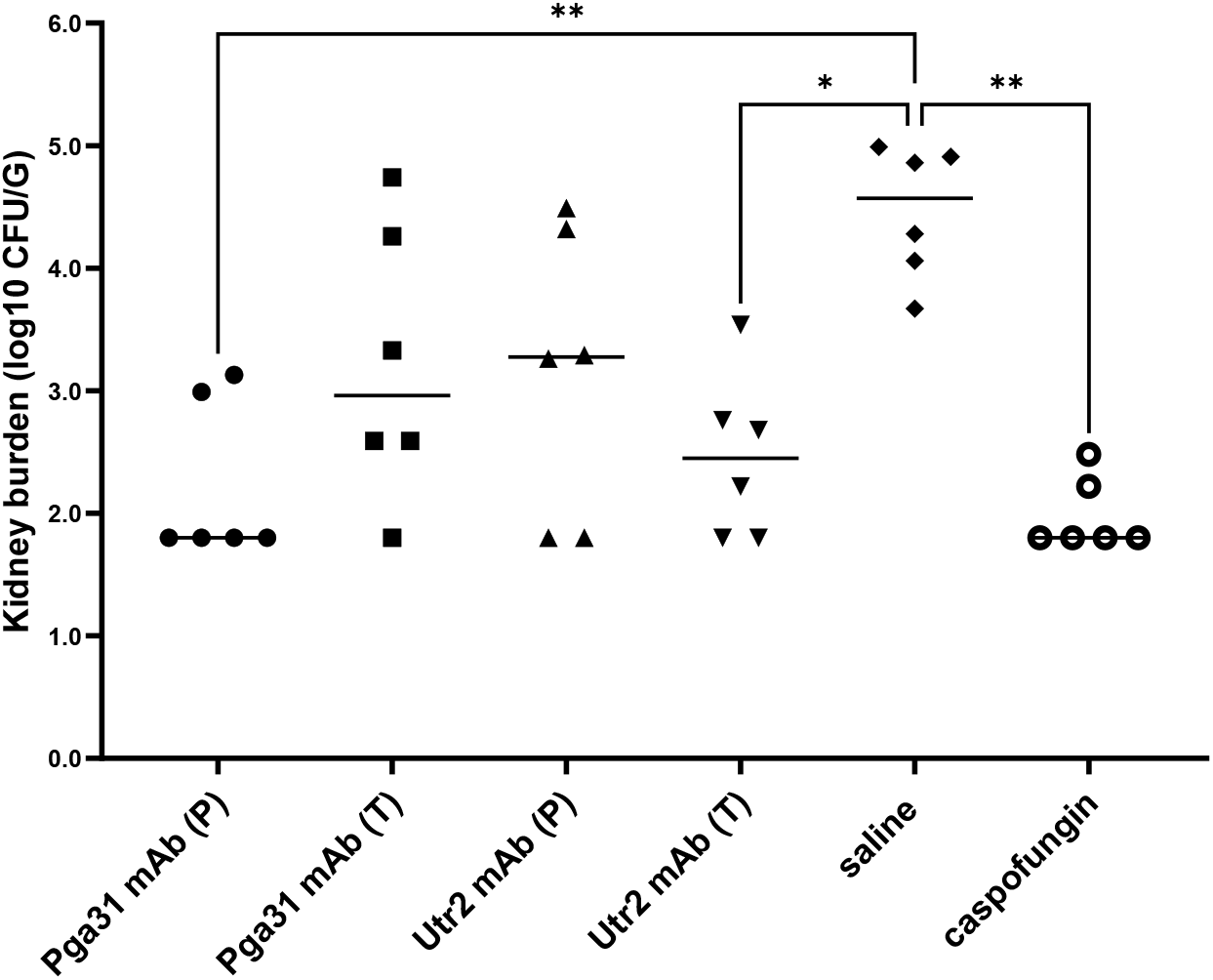
Kidney burdens at day 7 post-infection in prophylactic vs treatment study. Test mAbs (12.5 mg/kg per mouse) were administered 3 h pre-infection and/or 24 h and 72 h post-infection (n= 6 mice/group). Pga31 mAb (P): prophylactic arm, antibody given pre- and post-infection, Pga31 mAb (T): treatment arm, antibody given post-infection only, Utr2 mAb (P): prophylactic arm, Utr2 mAb (T): treatment arm, saline only control, caspofungin at 1 mg/kg body weight post-infection at 24 h and 72 h post-infection. Each symbol represents an individual mouse and bar represents mean kidney burden in each group. The detection limit for kidney burden determination was 2.3 log_10_ CFU/g and therefore any samples with zero count were assigned a value of one-half log below the detection limit (i.e. 1.8 log_10_ CFU/g). Kidney burdens for the different groups were compared by Kruskal-Wallis and Dunn’s multiple comparison tests, *P < 0.05, **P < 0.01, ***P < 0.005.

## DISCUSSION

With the advent of mAb technology, several groups have reported the development of protective antibodies as a central part of a patient’s recovery from infection (31). These mAbs typically recognise antigens that are unique to the fungus and include fungal cell wall polysaccharides and a small number of cell wall proteins (Als3, Sap2, Hsp90 and Hry1) involved in cell growth, virulence and pathogenesis, as reviewed in (15) (32). Utr2 and Pga31 are CWPs covalently linked to the fungal cell wall with their level of expression affected by carbon source (33) and to infection-associated stress-conditions, including external stimuli such as challenge with antifungal agents (34). We have recently reported an increased expression of Utr2 and Pga31 proteins, at proteomic levels, in *C. albicans* grown in the presence of caspofungin (22). In most cases, the C terminal end of GPI-anchored proteins are buried inside the β-glucan skeletal layer with only stretches of amino acids at the N terminal functional domain being surface exposed (35). Surface epitopes of Utr2 and Pga31 were deduced from tryptic peptides generated from our cell wall proteome studies and recombinant mAbs isolated that recognised CWPs in their native conformation (Fig. 3D-F). Pga31 antibodies showed increased binding to *C. albicans* whole cell lysates grown in the presence of caspofungin (Fig 1C-D), reaffirming observations that antifungal agents alter CWP expression (22) and that Pga31 is expressed as a remodelling mechanism for maintaining wall integrity under cellular stress. A role for Utr2 in establishing a compensatory mechanism for the crosslinking of chitin and β 1-3 glucan in echinocandin-treated cells has been reported previously (18). The abundance of Utr2 protein in the wall of caspofungin-treated cells is confirmed here, with increased antibody binding to cell lysate preparations (Fig. 2D-F). The target specificity of these antibodies was confirmed by a lack of binding to mutant strains (*utr*2Δ and *utr*2Δ/*crh11*Δ/*crh12*Δ), even in the presence of caspofungin (Fig. 2G).

We observed *C. albicans* cell morphology dependant immunoreactivity of CWP antibodies, with preferential binding to the hyphal form when compared to yeast cells. Utr2 mAb 1D2 bound uniformly along the growing hyphae, indicating broad surface exposure of this antibody’s preferred epitope. In contrast, the second Utr2 mAb 1H3, displayed a different binding pattern, mostly localised to the apical tip of growing hyphae suggesting recognition of a second and distinct epitope at the tip of the germ tube during hyphal elongation. The significance of the Crh family of proteins including Utr2 in cell wall biogenesis and their temporal and spatial organisation in various morphologies has been elegantly reported previously (18). This current study supports the idea that Utr2 accumulation initially is localised to new budding sites in yeast during the early growth phase, followed by re-localisation towards the base of the bud neck, overlapping with the chitin ring later in the cell cycle. 1H3 and 1D2 mAb binding to cells pre-treated with caspofungin also saw the greatest signal intensity at the new bud surface, further supporting this finding.

Pga31 mAb binding resulted in a weaker but punctate signal in distinct hyphal regions of *C. albicans* cells. This binding pattern is in agreement with previous reports and our own finding that Pga31 is expressed at low or undetectable levels in *C. albicans* under normal laboratory growth conditions. Pga31 is suggested to be part of the cell salvage pathway (16) and regulates chitin assembly when cells are treated with cell wall perturbing agents, including calcofluor white and caspofungin (12). In our study a marked increase in Pga31 binding was also observed when cells were treated with caspofungin (Fig 4B).

The ability of J774.1 macrophages to engulf *C. albicans* cells treated with CWP specific mAbs or an isotype control anti-*Candida* mAb was significantly higher than non-antibody treated cells (Table 2). A complex interplay between macrophage and *C. albicans* has been previously reported, with the pathogen sometimes counteracting the macrophage’s defence strategies to eventually break free and kill the macrophage as it escapes (36). In our study, whilst the antibody mediated engulfment did not result in complete killing of *C. albicans*, a significant inhibition of hyphal filamentation was observed which is tempting to speculate is due in part to an immunomodulatory activity of mAbs involved in pathogen clearance. This was further verified in a disseminated candidiasis mouse model, where protection from a life-threatening infection was evident in animals receiving CWP-specific mAbs compared to an isotype control mAb or vehicle alone. The Pga31 mAb, in particular, when administered as a single dose pre-infection, followed by two doses at 24 h and 72 h post-infection, conferred improved survival (compared to a single dose pre- and post-infection) of 83% vs 66%, respectively. The benefit of double dosing was also reflected in the kidney fungal burdens with a very respectable three log reduction (99.9%) in the number of fungal cells seen in the kidneys of mice receiving two doses of mAb post-infection.

Other published *in vivo* efficacy models typically describe test mAbs that were either pre-incubated with *C. albicans* cells or administrated as a prophylactic pre-pathogen challenge, with survival benefit and any reduction of fungal burden in associated organs reported (37) (27). With these experimental design parameters, mAbs are already present in the systemic circulation and able to bind to yeast cells, mediating opsonophagocytosis and clearance with enhanced protection. Survival rates between 40% and 50% in a mouse model of systemic candidiasis have been claimed for the β-(1→3)-D-glucan mAbs (37) and, in a separate study, less than one log reduction in kidney fungal burdens was reported for an anti-*C. albicans* mAb isolated from patient B cells (27). Using a peptide biologic therapy, rather than a much larger antibody, a small protective effect in animal studies with a one log drop in fungal burden was only seen in “topical” vaginal and oropharyngeal candidiasis models (38). This less potent systemic efficacy may be the result of the peptides “sticky” mode of action, low bioavailability or significantly reduced half-life compared to mAbs (39) (40). Interestingly, in our study 3 design, CWP-specific antibodies also reduced the fungal burdens in the kidneys of mice receiving treatment 24 h post infection, providing an early indication of their ability to bind *in vivo* to both yeast and hyphal morphologies, possibly inhibiting cell replication and/or enhancing phagocytosis and clearance. In the case of the anti-Utr2 mAb, double dose treatment did not translate into increased survival compared to the single dose (33% in both study 1 and 2). But in study 3, specifically investigating kidney fungal burdens, two doses of Utr2 mAb in the treatment only group was more protective than mice receiving mAb as a prophylactic, followed by two doses post infection. This data appears to reinforce our earlier observation that Utr2 mAb preferentially binds to the hyphal form of *C. albicans* (Fig 4A).

With the serum half-lives of therapeutic human IgGs often reported in the region of 21-28 days (a few days in mice) (41), immunotherapy for invasive fungal infections can reduce the dosing frequency whilst addressing the serious drug resistance issues associated with long term treatment regimens adopted in chronic infections. This is particularly pertinent for non-*albicans* species including *Candida krusei*, *C. glabrata* and *C. auris* which are either intrinsically tolerant or fully resistant to one or more class of existing antifungals, increasing the prevalence of non-treatable nosocomial infections (42). Off-target toxicity and drug-drug interactions are other important treatment considerations associated with existing antifungal therapies. Each drug class has its own clinical challenges: nephrotoxicity of polyenes (43), amphotericin B interactions with hypokalaemic drugs resulting in cardiac and skeletal muscle toxicity, and azole-mediated inhibition of metabolising liver enzymes (e.g. cytochrome P450) leading to decreased catabolism of co-administered drugs and dose related toxicities (44). It is widely recognised that targeted biological agents, such as monoclonal antibodies, can overcome many of the toxicity and drug resistance hurdles created by using small molecule drugs, with a handful of mAbs now approved by the FDA as prophylactic treatments against bacterial and viral infections (45). One could easily envisage scenarios where patients undergoing chemotherapy or organ transplant might receive long-acting, antifungal mAbs as prophylaxis against life-threatening invasive fungal infections prevalent in these immuno-compromised groups. These mAbs could be used as part of a co-therapy regimen to augment or prolong the activity of existing, but limited in number, systemic antifungals and address the issue of drug resistance, especially with fungistatic azole compounds. By simultaneously targeting the pathogen with an antifungal agent and a mAb adjuvant, the same level of efficacy can be achieved with lower drug doses, thereby significantly improving the narrow therapeutic window associated with commonly used antifungals.

In this present study we successfully identified mAbs as lead molecules in a new antifungal drug class as the first step in providing alternatives to our current and very limited antifungal drug portfolio. A panel of recombinant, fully-human, monoclonal antibodies to surface exposed epitopes of key fungal cell wall proteins in a range of fungal pathogens have been isolated and characterised. These antibodies have been selected to recognise target proteins in their native conformation and confer excellent levels of protection (over 80%) in a mouse model of disseminated candidiasis, in part, through the recruitment of phagocytic macrophages via antibody mediated opsonisation. These novel antifungal mAbs are now entering later-stage preclinical evaluation to investigate additional aspects of their mode of action, levels of *in vivo* tolerability and pharmacological activities including PK/PD profiles, in relevant animal models.

## MATERIALS AND METHODS

### Fungal strains, media and growth conditions

All fungal strains used in this study are shown in Table 4 and were cultured from glycerol stocks (−70 °C) and maintained on YPD agar plates containing 2% (w/v) glucose, 2% (w/v) mycological peptone, 1% (w/v) yeast extract, and 2% (w/v) agar (all from Oxoid, Cambridge, UK). For *C. albicans* strains, unless stated otherwise, a single colony was grown in YPD medium (see above without the agar) and grown overnight at 30 °C with shaking at 200 rpm. To induce hyphal formation, cells were grown in RPMI-1640 modified medium (Sigma-Aldrich) containing 10% heat-inactivated foetal calf serum (FCS) and incubated at 37 °C for 2-4 h. For mouse studies, *C. albicans* SC5314 was grown in NGY medium containing 0.1% Neopeptone (BD, Wokingham, UK), 0.4% glucose, 0.1% yeast extract (BD) at 30 °C with constant rotation at 200 rpm.

**Table 4.** C. albicans strains used in this study.

| Strain | Description | Genotype | Reference |
| --- | --- | --- | --- |
| SC5314 | Clinical isolate | Wild-type | (46) |
| GPY03 | *utr2Δ* | *ura3Δ::λimm434/ura3Δ::λimm434, utr2Δ::hisG/utr2Δ::hisG-URA3-hisG* | (18) |
| GPY102 | *crh11/crh12/utr2Δ* | *ura3Δ::λimm434/ura3Δ::λimm434, utr2Δ::hisG/utr2Δ::hisG; crh12Δ::hisG/crh12Δ::hisG; crh11Δ::hisG crh11Δ::hisG-URA3-hisG;* | (18) |
| CAMY 204 | *UTR* overexpression strain | *ura3Δ::λimm434/ura3Δ::λimm434, his1Δ::hisG/HIS1, arg4Δ::hisG/ARG4 ADH1/ADH1/adh1::P<sub>ADH1</sub>-cartTA::SAT1, RPS1/rps1::Clp10 - GTW- P<sub>TET</sub>-UTR2* | This study |
| *pga31Δ* | *pga31Δ* | *ura3Δ::λimm434/ura3Δ::λimm434,*<br>*arg4::hisG/arg4::hisG,*<br>*his1::hisG/his1::hisG*<br>*pga31::URA3/pga31::ARG4,* | (12) |
| CAM_K46 | *PGA31*<br>overexpression<br>strain | *ura3Δ::λimm434/ura3Δ::λimm434,*<br>*his1Δ::hisG/HIS1, arg4Δ::hisG/ARG4*<br>*ADH1/adh1:: P<sub>ADH1</sub>-cartTA::SAT1,*<br>*RPS1/rps1::Clp10 -GTW- P<sub>TET</sub>-PGA31* | This study |

### Phage display based isolation of CWP scFv binders from a human antibody library

Solution phase biopanning and monoclonal phage binding enzyme-linked immunosorbent assay (ELISA) were performed according to previously published methods (47). Briefly, a human antibody library was subjected to repeated rounds of selection using biotinylated peptide antigens corresponding to surface exposed regions of cell wall proteins Pga31 and Utr2. In the first round, streptavidin magnetic beads (Dynabeads M280, Invitrogen) were coated with 500 nM of biotinylated Utr2 or Pga31 peptide and phage particles displaying antibody fragments on their surface were incubated for target binding. Phage particles bound to antigen-biotin complex were eluted by triethylamine (TEA) and amplified by infecting *Escherichia coli* TG1 cells. For the second and third rounds of panning, the coating concentration of biotinylated peptide antigen was reduced to 100 nM and 10 nM respectively and rescued phage from previous rounds of panning allowed to bind to the antigen for selection. For screening antigen specific phage binders, 96-well plates (Nunc Maxisorp) were precoated with streptavidin to capture biotinylated Pga31 or Utr2 peptides and monoclonal phage supernatant was added as described (48). Peptide antigen binding ELISA was performed and individual phage monoclonals specifically binding to CWP peptide antigen (and not recognising non-related biotinylated peptide) were selected for antibody gene sequencing.

### Expression of soluble Pga31 and Utr2 antibody fragments (scAbs) in a bacterial system

Positive phage clones with unique VH and VL genes were converted into soluble single chain antibodies (scAbs) by cloning their respective scFv gene (VH-linker-VL) into the bacterial expression vector pIMS147 and transforming *E. coli* TG1 cells for periplasmic expression as described (30). ScAbs expressed in the bacterial periplasm were released by adding osmotic shock solution consisting of Tris-HCl-sucrose and EDTA followed by MgSO_4_ and incubated on ice, gently shaking for 15 min each. Recombinant scAbs were purified using immobilized metal affinity chromatography (IMAC) columns via binding of hexa histidine tagged protein to activated Ni-sepharose beads and elution using imidazole. Purified scAbs were dialysed against PBS and quantified either by SDS-PAGE, where the intensities of protein bands were compared (Image J), or by calculating final scAb concentrations by measuring the absorbance values at 280 nm using Ultraspec 6300 pro UV/Visible spectrophotometer (Amersham, Biosciences).

### Reformatting CWP scAbs into human-mouse chimeric mAbs

Utr2 and Pga31 scAbs were reformatted into human-mouse (IgG2a) chimeric mAbs by inserting the antibody VH and Vκ genes into a dual plasmid eukaryotic vector system encoding constant heavy and light chain genes of mouse IgG2a isotype. VH and Vκ genes of shortlisted scAbs were custom synthesised by introducing the restriction sites BssHII and BstEII (for VH gene) and BssHII and XhoI (for Vκ gene) at their 5ˈ and 3’ ends respectively (GeneArt custom gene synthesis service by Thermofisher), cloned into respective eukaryotic expression vectors pEEDM2a (encoding mouse IgG2a constant regions) or pEEDMκ (for mouse κ constant domain) using standard restriction enzyme digestion and ligation steps. Ligated DNA was purified using ethanol precipitation and used to transform electrocompetent *E. coli* TG1 cells for plasmid propagation.

### Production of Pga31 and Utr2 mAbs in a mammalian expression system

For laboratory scale expression of mAbs, plasmids bearing chimeric antibody heavy and light chain genes were prepared (EndoFree Plasmid Mega prep Kit, QIAGEN) and transfected into Human Embryonic Kidney cells (HEK293F) (Life Technologies) using polyethylenimine (PEI). Transfections were carried out using 1 mg of total DNA (500 μg each of VH and Vκ plasmid DNA) and 1 l of cultured HEK293-F cell suspension maintained in sterile Freestyle 293 expression medium (Invitrogen) without antibiotics at 37 °C, with 8% CO_2_, 125 rpm shaking. The transfected cells were grown for 8 days and purified using ProSep A beads (Millipore) and Econo-Pac chromatography columns (Biorad). Recombinant mAbs were eluted in 100 mM glycine (pH 3.0) before neutralisation with 1 M Tris-HCl (pH 8.0). Purified mAbs were quantified by SDS PAGE and A280 nm measurements.

### CWP Peptide, *C. albicans* cell lysate, and whole cell ELISA

For ELISA using Pga31 or Utr2 peptides, 96 well Nunc Maxisorp plates were pre-coated with 100 nM streptavidin and blocked with 2% Marvel in PBS before adding 500 nM biotinylated peptides. ScAb or mAb samples were incubated with the antigen in doubling dilutions and the binding was detected using anti-Human C Kappa horseradish peroxidase (HRP) conjugated secondary antibody (Sigma) (for scAbs) or anti-Mouse IgG (H+L) HRP secondary antibody (Thermo Scientific) (for human-mouse chimeric mAbs).

For whole yeast cell or total cell lysate ELISA using wt, *pga31*Δ, *crh11/crh12/utr2*Δ and *PGA31* overexpression strains of *C. albicans*, overnight cultures were inoculated into fresh YPD medium with a starting OD_600 nm_ per ml of 0.1-0.2, grown at 30 °C until OD_600 nm_ per ml = 0.5-0.6 was reached, then caspofungin (0.032 μg/ml) was added for 90 min.

For total cell lysate preparation, caspofungin treated or non-treated cells were harvested and centrifuged for 5 min at 4000 rpm, washed with sterile dH_2_0 and 10 mM Tris-HCl pH 7.5 before resuspending again in fresh Tris-HCl. Glass beads (0.5 mm diameter) were added (0.5 g to each 100 mg pellet) along with protease inhibitor solution (complete mini EDTA-free protease inhibitor cocktail, Roche), dH_2_O and 1 mM phenylmethylsulfonyl fluoride (PMSF) in ethanol. Samples were subjected to 15 rounds of bead beating for 35 sec at speed 6.5 using a Fast Prep^®^-24 instrument (MP Biomedicals, UK) with tubes placed on ice for at least 5 min in between each round of bead beating. After centrifugation at 3000 rpm for 1 min to pellet the beads, the broken cells suspension was transferred to sterile cold tubes. Cell lysate preparation was used to coat ELISA plates as before.

For ELISA using *Candida* hyphae, cells were grown in RPMI-1640 medium (for 2-4 h) and added to the wells for incubation at 37 °C for 1 h. For whole cell binding ELISA, *Candida* coated wells were washed and blocked with 2% BSA (Sigma), followed by the addition of double diluted scAb or mAb samples. Binding was detected using anti-Human C Kappa HRP or anti-Mouse IgG (H+L) HRP and the resulting immunoreaction was measured as described previously.

### Immunofluorescence imaging of antibodies binding to fungal cells

Fungal cultures, grown as described above, were diluted 1:1000 in milliQ water and left to adhere on a poly-L-lysine glass slide (Thermo Scientific, Menzel-Gläser) for 1 h. Cells were washed three times in Dulbecco’s phosphate-buffered saline (DPBS) and fixed with 4% paraformaldehyde at room temperature. Blocking was done using 1.5% BSA which was followed by washing and cell staining using mAbs at 25 µg/ml for 1 h at room temperature. Alexa Fluor® 488 goat anti-mouse IgG antibody (Life Technologies) at 1 in 1000 dilution was added to the slide for 1 h at room temperature and washed prior to staining with 25 µg/ml calcofluor white (CFW) to stain cell wall chitin. Mounting medium and a coverslip were added before images were taken using an UltraVIEW® VoX spinning disk confocal microscope (Perkin Elmer, Waltham, Mass, USA).

### *Ex vivo* Macrophage Interaction assay

#### Macrophage culture

J774.1 mouse macrophage-like cells (ECACC, Salisbury, UK) were cultured in Dulbecco’s Modified Eagle Medium (DMEM, Thermo Fisher) supplemented with 200 U/ml penicillin/streptomycin, 2 mM L-glutamine (Invitrogen), and 10% (v/v) heat-inactivated foetal calf serum in tissue culture flasks at 37 °C with 5% CO_2_.

For interaction assays, macrophages were seeded at a density of 1 × 10^5^ cells per well of a glass imaging dish (Ibidi) and incubated overnight at 37°C, 5% CO_2_. Immediately before phagocytosis experiments, supplemented DMEM was replaced with pre-warmed supplemented CO_2_-independent medium (Thermo Fisher) to ensure macrophages remained viable during the analysis of *C. albicans* interactions.

#### *C. albicans* preincubation with test antibodies

Prior to phagocytosis assays, *C. albicans* SC5314 yeast cells (3 × 10^5^ cells) were either untreated or pre-coated with 50 µg/ml anti-Pga31, anti-Utr2 mAbs or a commercially sourced anti-*Candida* mouse IgG2a monoclonal antibody (MA1-7009) (Fisher Scientific) in pre-warmed supplemented CO_2_-independent medium and incubated at 37 °C with gentle shaking for 40 min.

#### Live cell video microscopy of phagocytosis assay

Video microscopy experiments were performed using an UltraVIEW® VoX spinning disk confocal microscope in a 37°C chamber, with images captured at 1 min intervals over a 2 h period using a Nikon camera (Surrey, UK). Six different videos were recorded for each antibody or control group from two biological replicates, and subsequent analysis was conducted using Volocity 6.3 imaging analysis software (PerkinElmer).

At least 25 macrophages were selected at random from each video to determine phagocytic activity. Measurements taken included: (a) time of engulfment, defined as time between macrophage establishing cell-cell contact to complete engulfment of the *C. albicans* cell and (b) the length of intracellular hyphae at two time points (60 and 90 min). Mean values of the length of 25 intracellular hyphae for each time point were calculated. A Shapiro-Wilk test for normality was used to determine the distribution of data where appropriate. Kruskal–Wallis test with Dunn’s multiple comparisons test was used to determine statistical significance using GraphPad Prism 5.

### Investigating the therapeutic efficacy of CWP mAbs in mouse disseminated candidiasis model

All animal experimentation was done in accordance with UK Home Office regulations and was approved by both the UK Home Office and an institutional animal welfare and ethical review committee (AWERB). Female BALB/c mice, 7-9 weeks old (Envigo, Huntingdon, UK) were randomly assigned to groups of 6 for treatment and control. *C. albicans* inoculum was prepared by growing strain SC5314 in NGY medium for 16 h, with shaking at 30°C. Cells were harvested and washed with saline, counted by haemocytometer, and resuspended in saline to provide an inoculum of approx. 2 x 10^4^ CFU/g mouse body weight in 100 µl. Mice were infected intravenously, and the actual inoculum level determined by viable cell counts. Depending on the study, the treatment dose of mAbs was either 12.5 mg/kg or 15 mg/kg per mouse in 150 µl. MAbs were administered as prophylaxis or treatment. In prophylactic studies, a single dose of antibody was delivered 3 h prior to challenge with *Candida* inoculum followed by either single mAb dosing 24 h post infection and double dosing 24 and 72 h post infection. For treatment only group, two doses of mAb were administered 24 and 72 h post *Candida* challenge. The comparator drug caspofungin was dosed at 1 mg/kg at 24 h and 72 h post-infection. Vehicle only control followed the same dosing pattern of test mAbs.

Mice were monitored and weighed every day during the course of experiments and at the end of study period, mice were culled by cervical dislocation and organs, including kidneys, spleen and brain, were removed aseptically and used to determine fungal burdens by plating out organ homogenates and counting colonies after 24 h growth at 30°C.

In survival studies, mice which lost more than 20% of their initial body weight and/or showed signs of a progressive systemic infection were culled by cervical dislocation and their day of death recorded as occurring on the following day.

### Statistical Analysis

Statistical analyses of mouse survival data were carried out with IBM SPSS and GraphPad Prism 5 was used for rest of the data. For antibody binding curves, data points are expressed as mean ± SEM (n=2). For macrophage assay and fungal burden in mouse kidneys and other organs, results are shown as mean ± SD. When comparing two or more groups, a Kruskal–Wallis test with Dunn’s multiple comparisons was performed to determine statistical significance, across all groups, then between different groups when there was a difference across all groups. Mouse survival estimates were compared by the Kaplan-Meier log-rank test.

## Acknowledgments

The authors gratefully acknowledge Kevin McKenzie and Lucy Wight from the University of Aberdeen Microscopy and Histology Facility for training and access to fluorescence microscopy and for their support & assistance in this work. The authors also gratefully acknowledge Dr David Stead from Aberdeen Proteomics for his support & assistance with the *Candida* proteome analysis and the staff of the University of Aberdeen Medical Research Facility for their support & assistance with the mouse studies.

## Funding

This work was supported by the following research grants:

High throughput and Fragment screening Fund, Scottish Universities Life Sciences Alliance (SULSA)

Seed Corn Award from the University of Aberdeen Wellcome Trust Institutional Strategic Support Fund

MRes Studentship by the Medical Research Council Centre for Medical Mycology at The University of Aberdeen (grant number MR/P501955/1)

PhD studentship, Institute of Medical Sciences, University of Aberdeen

PhD studentship, Taibah University and Saudi Government Scholarship

PhD studentship by the European Union’s Horizon 2020 research and innovation program under the Marie Sklodowska-Curie grant agreement no: H2020-MSCA-ITN-2014-642095 (OPATHY)

## Author contributions

CAM, SP, AJP contributed to the concept and study design, CAM and SP developed the methodology. SAA and LF performed recombinant antibody generation and ELISAs, LF and MW completed IgG reformatting and production of mAbs for animal studies; MM, THT and LAW performed ELISAs and IF staining; MM performed macrophage assays and DM planned, conducted and analysed animal studies. CAM, SP, AJP contributed towards funding acquisition and project administration and CAM, SP and LAW towards the supervision and training of MRes and PhD students. SP wrote the original draft and CAM, DMM and AJP completed review and editing. All authors had full access to the data and approved the manuscript before it was submitted by the corresponding author

## Data and materials availability

All data are available in the main text or the supplementary materials. Antibodies described in this paper will be available for research purposes through a material transfer agreement with the University of Aberdeen

## Supplementary Materials

**Fig. S1.**
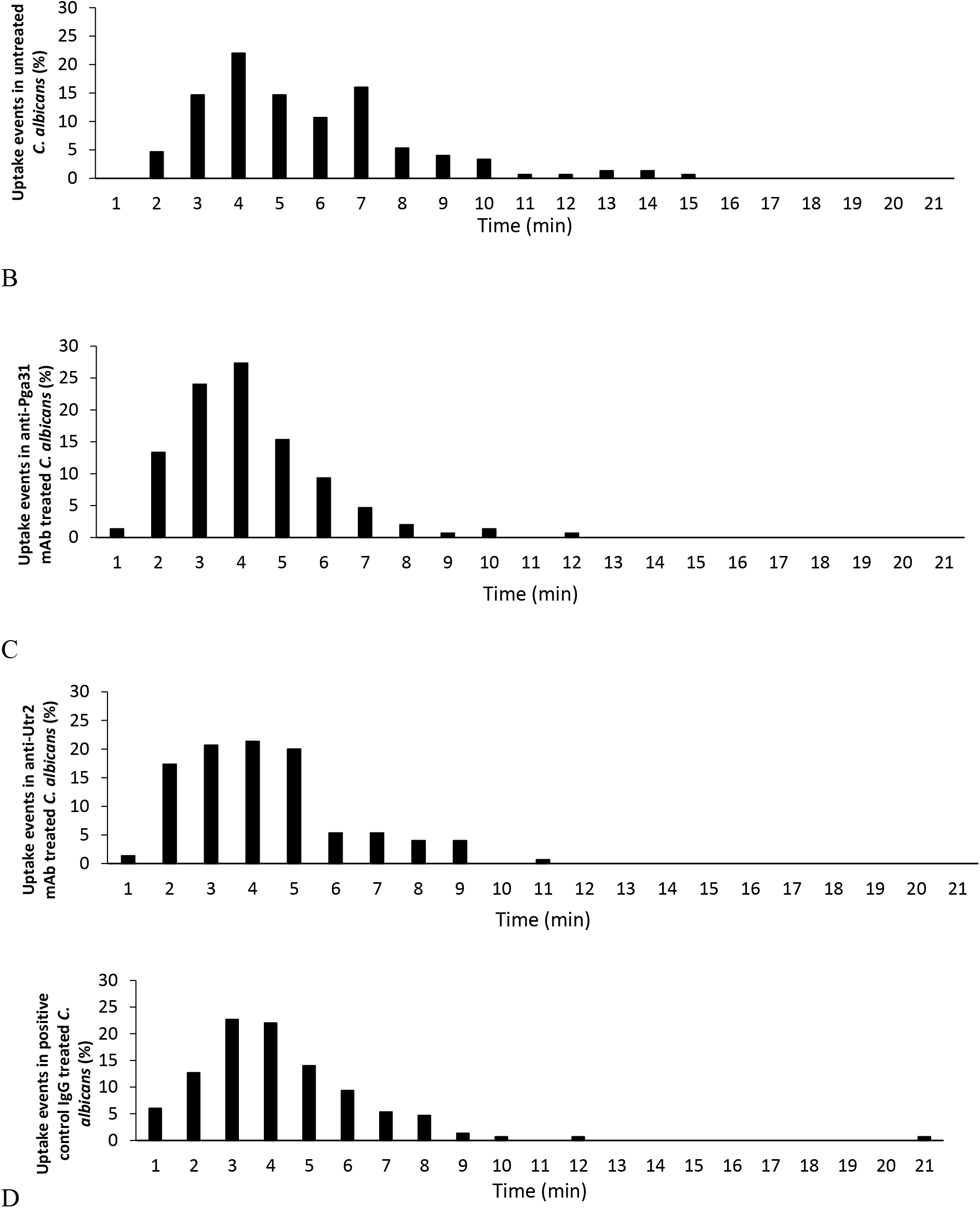
Engulfment of anti-Candida mAb treated C. albicans cells by mouse macrophages. The time taken for J774.1 macrophages to ingest live Candida cells following initial cell-cell contact verses the percentage of uptake events are plotted. (A) wt (B) Pga31 mAb (C) Utr2 mAb (D) positive control murine mAb. The rate of engulfment of all antibody-treated cells was faster than that of untreated *C. albicans*. Bars represent the percentages of uptake events (n = 6 videos for each antibody group from two biological replicates).

**Table S1.**
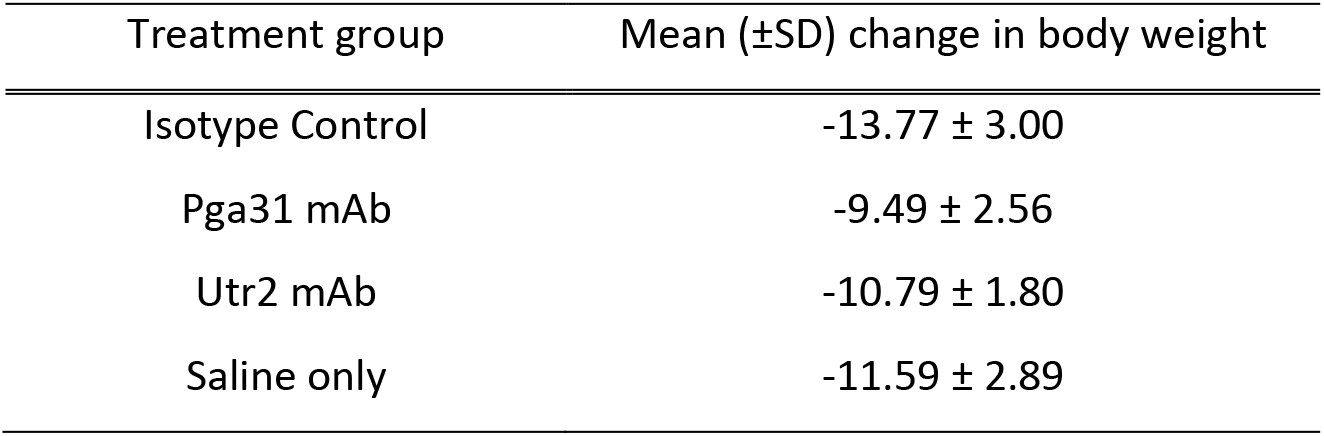
Average weight change in mice in study 1. Groups of mice (n=6) were treated with either Pga31 mAb (15 mg/kg), Utr2 mAb (15 mg/kg), mouse IgG2a isotype control (15 mg/kg) or saline, 3 h pre and 24 h post-infection in a murine model of disseminated candidiasis. Data represents mean change in body weight ± SD (g) at day 2 compared with day 0 in mouse study 1.

